# Novel imprints in mouse blastocysts are predominantly DNA methylation independent

**DOI:** 10.1101/2020.11.03.366948

**Authors:** Laura Santini, Florian Halbritter, Fabian Titz-Teixeira, Toru Suzuki, Maki Asami, Julia Ramesmayer, Xiaoyan Ma, Andreas Lackner, Nick Warr, Florian Pauler, Simon Hippenmeyer, Ernest Laue, Matthias Farlik, Christoph Bock, Andreas Beyer, Anthony C. F. Perry, Martin Leeb

## Abstract

In mammals, chromatin marks at imprinted genes are asymmetrically inherited to control parentally-biased gene expression. This control is thought predominantly to involve parent-specific differentially methylated regions (DMR) in genomic DNA. However, neither parent-of-origin-specific transcription nor DMRs have been comprehensively mapped. We here address this by integrating transcriptomic and epigenomic approaches in mouse preimplantation embryos (blastocysts). Transcriptome-analysis identified 71 genes expressed with previously unknown parent-of-origin-specific expression in blastocysts (nBiX: novel blastocyst-imprinted expression). Uniparental expression of nBiX genes disappeared soon after implantation. Micro-whole-genome bisulfite sequencing (μWGBS) of individual uniparental blastocysts detected 859 DMRs. Only 18% of nBiXs were associated with a DMR, whereas 60% were associated with parentally-biased H3K27me3. This suggests a major role for Polycomb-mediated imprinting in blastocysts. Five nBiX-clusters contained at least one known imprinted gene, and five novel clusters contained exclusively nBiX-genes. These data suggest a complex program of stage-specific imprinting involving different tiers of regulation.

## INTRODUCTION

Most mammalian genes are active on both parental alleles, but some are expressed from only one, determined by the parent-of-origin, and are said to be imprinted. Balanced expression of imprinted genes is critical ^1,2^ as development stops around the time of implantation in uniparental diploid embryos ^1,3,4^. Multiple databases of imprinted genes exist that collectively list 388 genes with parent-of-origin expression bias in mice ^5–7 1,8^ (http://www.geneimprint.com/). Imprinting is associated with chromatin marks that include allele-specific DNA methylation and/or H3K27me3 ^9^. DNA methylation-based imprints are associated with differentially methylated regions (DMRs). Many DMRs are established during gametogenesis in a Dnmt3l-dependent manner to produce germline DMRs (GL-DMRs) ^10^. GL-DMRs are key constituents of each of the 24 known imprinting control regions (ICRs) in the mouse ^6,11,12^.

Although uniparental embryos fail in early development, they form blastocysts from which pluripotent embryonic stem cells (ESCs) can be established. Haploid parthenogenetic (ph) and androgenetic (ah) embryos have been utilised for the derivation of haploid ESCs whose nuclei can be combined with complementary gametes to produce living mice, albeit inefficiently, in ‘semi-cloning’ ^13–16^. The extent to which poor development in semi-cloning reflects imprinting instability is unclear and the DNA methylomes of mouse uniparental embryos have not been comprehensively evaluated. However, it is known that haploid ESCs lose canonical imprints over extended culture periods, which has been leveraged to generate bi-maternal mice ^17^.

In addition to genomic DNA methylation-based imprinting, a subset of genes with paternal expression bias in mouse preimplantation embryos is maternally enriched for H3K27me3, without apparent direct dependence on DNA methylation. Most or all of this H3K27me3-based imprinting is lost in extra-embryonic cell lineages and post-implantation ^9,18^. However, the extent of imprinting control by both types of epigenetic mechanisms in mouse preimplantation development is unknown. Imprinting defects can have severe developmental consequences that can manifest themselves at, or shortly after implantation ^19^. It is therefore likely that the imprinting landscape in blastocysts is a critical determinant of normal development, such that blastocyst imprinting dysregulation has serious detrimental developmental consequences ^20^. We therefore sought to determine parent-of-origin-specific expression in biparental embryos and parent-of-origin-specific DNA methylation in uniparental blastocysts to delineate the imprinting landscape in mouse preimplantation embryos.

## RESULTS

### Assessing parent-of-origin-specific gene expression in F1-hybrid mouse blastocysts

To delineate parent-of-origin expression bias in mouse blastocysts, we performed allele-specific transcriptome analyses (RNA-seq) of embryonic day 3.5 (E3.5) embryos obtained without *in vitro* culture from reciprocal *Mus musculus domesticus* C57BL/6 (B6) x *Mus musculus castaneus* (*cast*) natural mating (Fig. 1a). After exclusion of transcripts encoded by the X-chromosome, 10,743 robustly expressed transcripts were identified that contained informative single nucleotide polymorphisms (SNPs) (≥ 12 reads in at least four out of eight samples). The list included 134 of the combined catalogue of 388 genes previously reported to be imprinted ^1,8^ and in repositories (Mousebook, Otago, Geneimprint, Wamidex) (referred to here as ‘published imprinted genes’). We further categorized the group of published imprinted genes into those listed in at least one repository (67 ‘repository imprints’) or in three or more repositories (30 ‘high confidence repository imprints’).

**Fig. 1.**
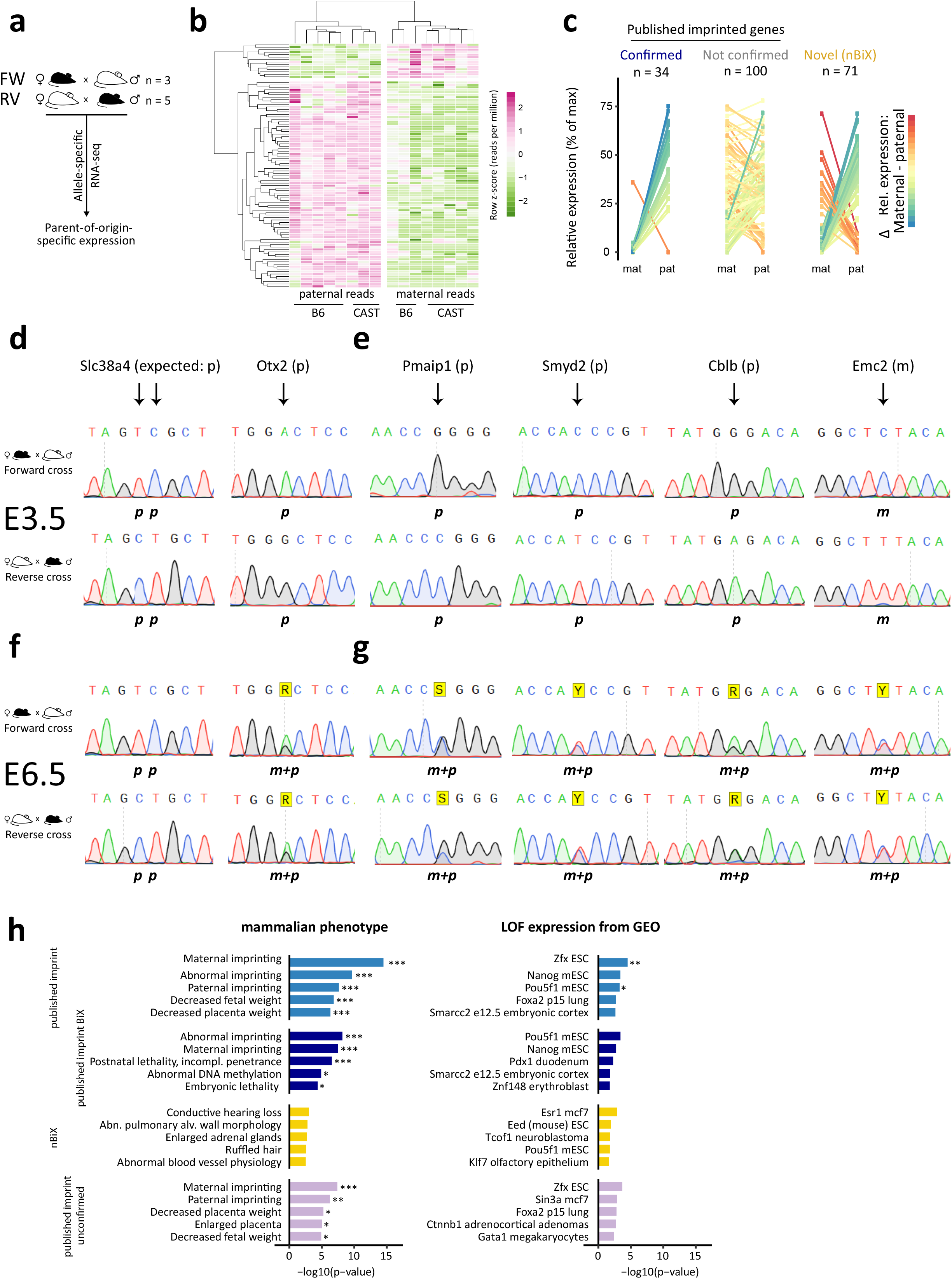
Parent-of-origin-specific gene expression in blastocysts. **a,** Schematic overview of experimental approach for the identification of parent-of-origin-specific gene expression. **b,** Heatmap showing row-normalized expression values of all 105 blastocyst imprint expressed (BiX) genes. Colour scale indicates Z scores based on reads per million. Maternal and paternal reads for the same sample are shown in separate columns. **c,** Distribution of SNP-containing RNA-seq reads between maternal and paternal alleles in different gene groups (confirmed published imprinted genes, unconfirmed published imprinted genes, novel imprinting candidate genes, nBiX). **d,** RT-PCR and Sanger sequencing-based validation of allele-specific expression of confirmed published imprinted genes *Slc38a4* and *Otx2* on embryonic day 3.5 (E3.5). **e,** RT-PCR and Sanger sequencing-based validation of allele-specific expression of nBiXs at E3.5. **f,** Sequencing electropherogram showing RT-PCR and Sanger sequencing-based validation of allele-specific expression of confirmed published imprinted genes *Slc38a4* and *Otx2* at E6.5. **g,** RT-PCR and Sanger sequencing-based validation of allele-specific expression of nBiXs at E6.5. **h,** Gene set enrichment analysis using Enrichr ^49^ of mammalian phenotypes associated with loss of confirmed, novel (nBiX), or unconfirmed parent-of-origin-specifically expressed genes (left). Enrichment analysis for the same gene sets compared to genes deregulated (either increased [up] or decreased [down] expression) in gene knockout and depletion experiments (right). The eight top terms based on p-value are shown for each category and odds ratios are plotted. *, adj . p < 0.05; **, adj. p < 0.01; ***, adj. p < 0.001.

Four hundred and nine (409) genes showed significant *cast*-specific, and 1001 significant B6-specific strain-dependent expression bias (adj. p ≤ 0.05, and further filtered for consistency across all samples). A further 141 genes exhibited parent-of-origin-specific expression in blastocysts (Fig. 1b; adj. *p* ≤ 0.1 and further filtered for consistency across all samples). To increase stringency, we imposed a requirement for consistent allelic expression ratio of at least 70:30 between parental alleles in a minimum of 60% of embryos for forward and reverse crosses. We refer to the first group of 141 transcripts as ‘blastocyst-skewed expressed’ genes (BsX), and those fulfilling the 70:30 criterion as ‘blastocyst-imprinted expressed’ genes (BiX; 105 transcripts).

BiX genes included 34 of the 134 genes from the combined catalogue of imprinted genes (35 for BsX; Fig. 1c and Extended Data Fig. 1a). Paternal expression of *Slc38a4*, *Peg3*, *Otx2* and *Bbx* was confirmed in independent reciprocal crosses by RT-PCR followed by Sanger sequencing (Fig. 1d; Extended Data Fig. 1c). A large proportion of the published imprinted genes (100 of 134, including *Igf2*, *H13* and *Commd1*) were absent from the BiX dataset (99 for BsX). We therefore re-evaluated whether these previously reported imprinted genes were indeed expressed equivalently from both alleles, or whether lack of statistical power may have prevented detection. To this end, we performed a statistical test for equivalent expression from paternal and maternal alleles. Across all analysed genes this identified statistically significant biallelic expression for 5376 genes (adj. *p* ≤ 0.1, H0: absolute log2FC ≥ 1), including 24 out of the 134 (18%) published imprinted genes with SNP-containing reads (Fig. 1c). RT-PCR Sanger sequencing of independent reciprocal cross E3.5 blastocysts revealed that *Commd1* indeed exhibited mixed expression states (with sample-dependent allele-specific or biallelic expression) and *Pon2* had clear strain-biased expression in blastocysts (Extended Data Fig. 1b). Hence, statistically significant parent-of-origin-specified gene expression was detected in blastocysts for only a quarter of all published imprinted genes. Of note, 18% of published imprinted genes were biallelically expressed in blastocysts with statistical significance, indicating a strong impact of tissue and cell type in defining imprinting patterns during development.

### Identification of novel imprinted genes

Groups of 71 (56 paternal, 15 maternal) and 106 (76 paternal, 30 maternal) genes that did not include published imprinted genes respectively constitute sets of novel BiX (nBiX) or novel BsX (nBsX) genes (Fig. 1c; Extended Data Fig. 1c). RT-PCR Sanger sequencing of independent crosses confirmed uniparental expression of the paternally-expressed nBiX genes *Pmaip1*, *Smyd2*, *Cblb*, *Myo1a* and of maternally-expressed nBiX *Emc2* in E3.5 blastocysts (Fig. 1e). Parent-of-origin-biased expression of tested nBiXs was lost by E6.5 (Fig. 1g), similar to the recently reported H3K27me3-dependent imprinted genes *Otx2* and *Bbx* ^9^, but in contrast to high confidence repository imprinted genes, *Slc38a4* and *Peg3,* which we found maintained uniparental expression at E6.5 (Fig. 1f; Extended Data Fig. 1e). We further confirmed parent-of-origin-specific expression of the BsX genes *Wrap53*, *Tmem144* and *Sri* (Extended Data Figure 1d and f) in independent crosses, suggesting consistent parental allele expression bias across multiple samples and experiments. Allele-specific expression analysis of available single-cell sequencing data confirmed parental expression bias of BiX and BsX (both novel and confirmed published imprinted genes), despite sparsity of signal (Extended Data Fig. 1g) ^18^. In accordance with our data, unconfirmed published imprinted genes did not exhibit clear allelic bias in the single-cell data. Our data accordingly identify sets of nBiX and nBsX genes with high confidence, parent-of-origin-specific blastocyst expression bias.

The nBiX set was enriched for genes up-regulated in cells lacking the PRC2 (Polycomb Repressive Complex 2) component *Eed* (Fig. 1h). *Eed* regulates H3K27me3 ^21^, a DNA methylation independent imprinting mark ^9,22,23^. Genes upregulated in ESCs upon loss of the pluripotency factors *Oct4* (*Pou5f1*) or *Nanog* were also enriched among nBiXs and confirmed published imprints, revealing an intersection between blastocyst imprinting and pluripotency circuitries. Both published confirmed imprinted and nBiX genes exhibited more dynamic transcriptional regulation during the first 24 hours of ESC differentiation ^24^ compared to biallelically-expressed genes (Extended Data Fig. 1h). This indicates that genes showing parent-of-origin-specific gene expression in blastocysts are involved in, or responsive to early embryonic cell fate specification.

### Capturing parent-of-origin-specific DNA methylation in uniparental mouse blastocysts

Imprinted gene expression has conventionally been associated with parent-of-origin-biased genomic DNA methylation. To assess whether parentally-specified nBiX expression could also be explained in this way, we measured genome-wide DNA methylation in individual haploid uniparental parthenogenetic haploid (ph) and androgenetic haploid (ah) E3.5 blastocysts by micro-whole-genome bisulfite sequencing (μWGBS) ^15,25,26^. We selected haploid embryos in an effort to reduce noise that might otherwise have been contributed by different alleles in diploid uniparental embryos. Moreover, uniparental embryos allow unambiguous mapping of μWGBS reads to chromosomes with known parental provenance. Uniparental embryos efficiently formed expanded blastocysts and contained cells expressing readily-detectable Oct4 and Sox2 (Extended Data Fig. 2a and b). For comparison, we also derived ahESC and phESC lines and included three androgenetic, four parthenogenetic and five biparental ESC lines and somatic tissue (kidney) in the μWGBS analysis (Fig. 2a).

**Fig. 2.**
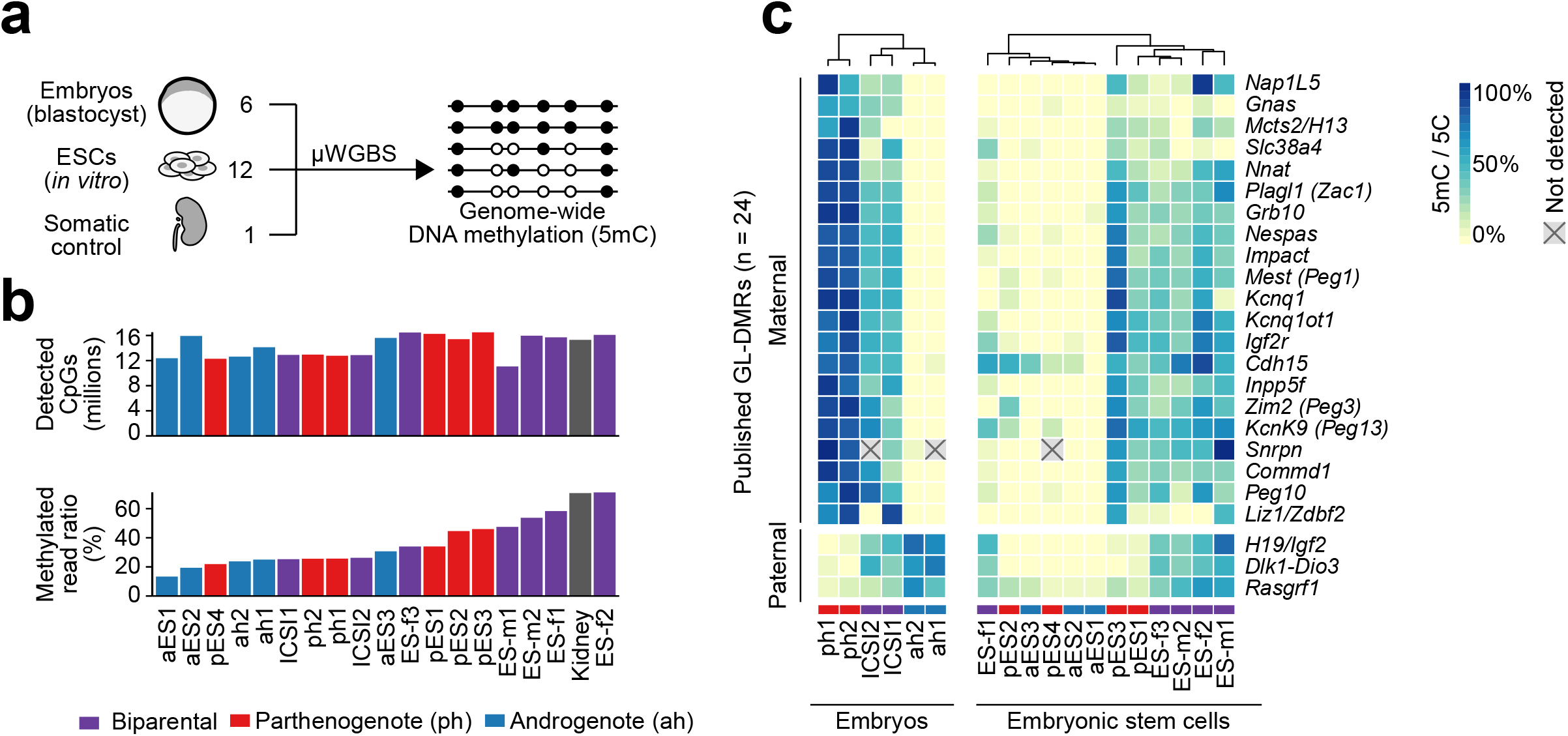
DNA methylation analysis confirms genomic imprints at known GL-DMRs. **a,** Schematic overview of samples used for DNA methylation analysis. **b,** Overview of the number of distinct CpGs detected and global DNA methylation levels in all samples. **c,** Heatmap showing DNA methylation levels (ratio of methylated out of all reads) in 24 known germline DMRs (GL-DMRs) in blastocyst samples (left) and ESCs (right).

DNA methylation levels could be quantified at 11 to 16.5 million CpGs per sample. Uniparental blastocysts exhibited ~25% global CpG methylation, independently of parental provenance, compared with ~70% CpG methylation in the kidney (Fig. 2b) and in line with 20% genomic methylation previously reported for the blastocyst inner cell mass ^27^. These data show that blastocyst global DNA methylation levels are independent of the parent-of-origin. Investigation of DNA methylation levels across published ICRs unambiguously confirmed parent-of-origin-specific methylation differences at most (22 out of 24) known GL-DMRs in haploid uniparental embryos (Fig. 2c). The two we did not detect across all samples were *Snrpn*, whose DMR lacked coverage in two samples, and *Liz1*/*Zdbf2*, whose DMR showed no evidence of DNA methylation in one of the intracytoplasmic sperm injection (ICSI)-derived replicates, confounding unambiguous identification. The uniparental embryo data thus efficiently detected GL-DMRs with a precision that may surpass that obtained for biparental F1 hybrid blastocysts or long read sequencing ^27,28^ (Extended Data Figure 2c).

Of six DMRs reportedly acquired in somatic tissues ^6^, we found that none were uniparentally methylated in blastocysts (Extended Data Figure 2d). The implication of this is that these loci carry marks at the blastocyst stage that are not directly dependent on DNA-methylation, and that do not become manifest as DNA methylation until after implantation. This is reminiscent of germ-line-independent somatic DMR acquisition on an *H19* transgene ^29^. The mechanism of this implicit DNA-methylation-independent preimplantation imprinting program and the means by which it is converted into DNA methylation, are unknown ^29,30^.

### DMR erosion in uniparental haploid ESCs

Four out of five biparental ESC lines maintained GL-DMR methylation levels similar to those in ICSI embryos when cultured in 2i/LIF medium; one female line (ESf1) exhibited erosion specifically of maternal DMRs (Fig. 2c), but imprint status was generally independent of sex and passage number. All biparental lines had strongly reduced DNA methylation of the *Gnas* ICR. The DNA methylation signal was reduced in some, but not all ESC lines on *Slc38a4* and *Liz1*/*Zdbf2* DMRs, suggesting differential stability of DMRs in ESC culture. In contrast, similarly-cultured haploid ESC lines typically underwent widespread DMR erosion regardless of their parental provenance. Androgenetic ESCs underwent near-complete erosion of all methylation over paternally-deposited *H19*/*Igf2* and *Dlk1* GL-DMRs (Fig. 2c). Methylation of the *Rasgrf1* GL-DMR was at lower levels than in biparental embryos, indicating ongoing loss of DNA methylation. Parthenogenetic haploid ESCs exhibited greater variability in GL-DMR methylation loss than their androgenetic (ahESC) counterparts. In two phESC lines, most maternal DMRs were maintained at levels similar to those of biparental ESCs, but even then, DMR signals were reduced compared to ph blastocysts, indicating that phESCs can maintain DMRs in culture, but undergo varying levels of DMR loss. Consistent with DMR dysregulation, ahESCs and phESCs lacked parent-of-origin biased expression of tested known imprinted genes (*Meg3*, *Peg3*, *Slc38a4* and *Jade1*), despite possessing unperturbed naïve pluripotency and the potential to initiate differentiation (Extended Data Fig.2e and f). Relative DMR stability in parthenogenotes compared to androgenotes contrasts with previous reports ^17^. These findings further support the idea that imprint erosion does not strictly reflect parental origin. However, imprint erosion in ahESCs would explain why the vast majority failed to support ‘semi-cloning’; embryos produced by injecting ahESC nuclei into mature oocytes would lack a balanced set of imprinted genes, resulting in developmental attenuation prior to, or around the time of implantation, as we observed ^3,4^. In summary, uniparental ESCs exhibited variable loss of DMRs, even though the DMRs were robustly detectable in uniparental blastocysts.

### Identification of novel blastocyst DMRs

Corroboration of known GL-DMRs allowed us to ask whether our data also revealed novel DMRs in blastocysts. Comparison of genome-wide DNA methylomes from ah and ph blastocysts with those of control blastocysts produced by intracytoplasmic sperm injection (ICSI) identified 859 DMRs (dmrseq, adj. p ≤ 0.1) ^31^. Of these, 778 (91%) were maternal (that is, the marks were enriched in parthenogenotes) and 81 (9%) paternal (enriched in androgenotes) (Fig. 3a). The DMRs were associated with 3,664 (7,031) and 392 (779) annotated genes within 100 (or 250) kb windows, respectively; 250 kb is well within the ~300 kb size of the *Igf2r* cluster ^2,32^. Unbiased embryo analysis recovered 23 of the 24 known GL-DMRs and in most cases coordinates of novel DMRs were superimposable upon those of published DMRs (Fig. 3b). For *Snrpn*, we detected a new DMR 1kb from the annotated *Snrpn* GL-DMR potentially extending the *Snrpn* DMR (Extended Data Fig.3a). Only the *Liz1/Zdbf2* GL-DMR was not confidently identified in our analysis, because it lacked a DNA methylation signal in one of the ICSI samples (Fig. 2c). As for known ICRs, patterns of blastocyst-derived DMRs identified here were not maintained in haploid ESCs (Extended Data Fig.3b).

**Fig. 3.**
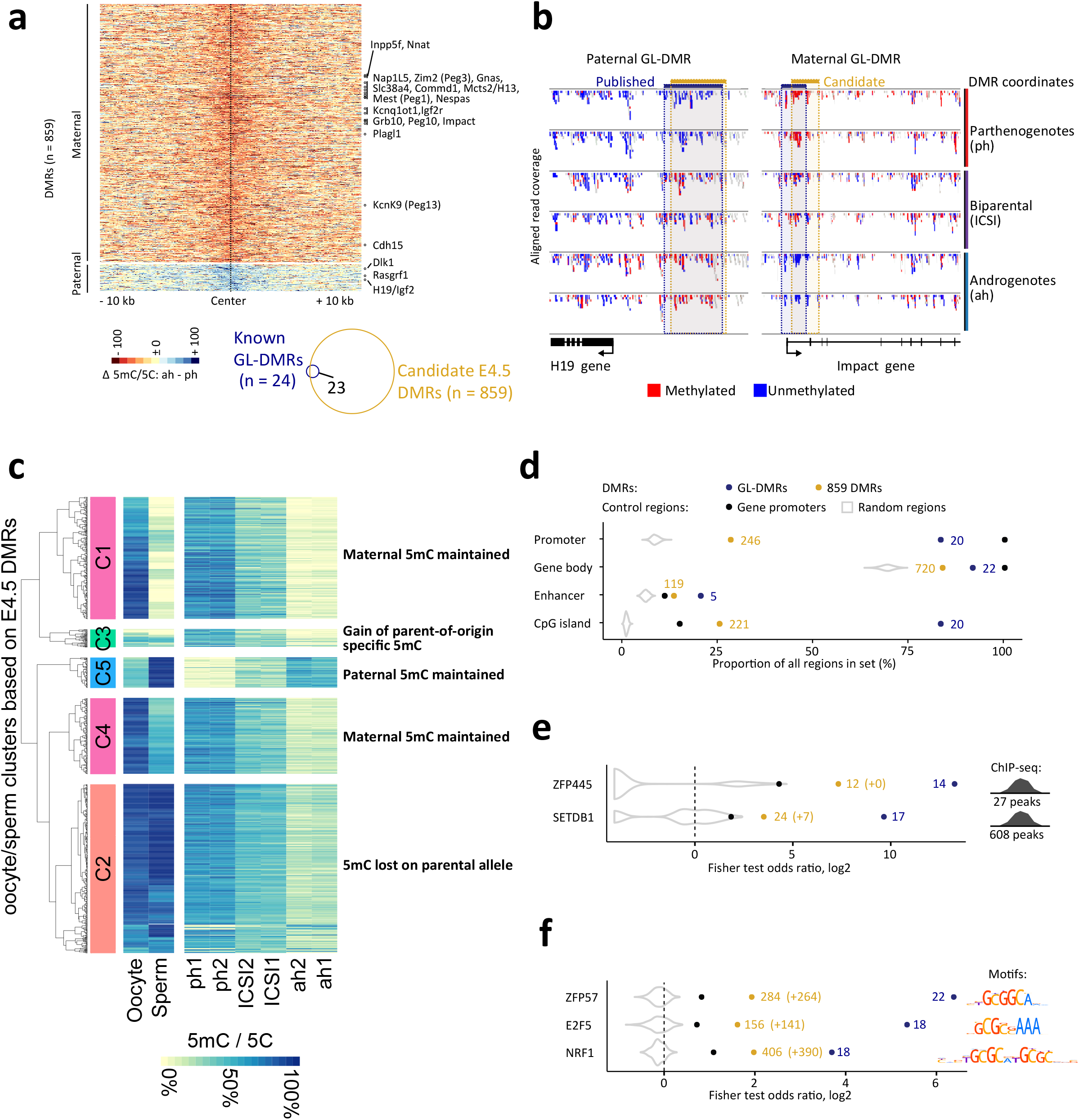
Novel DMRs in uniparental embryos. **a,** Heatmap showing DNA methylation signals over all 859 identified DMRs (red, maternal DNA methylation; blue, paternal DNA methylation). Known GL-DMRs are indicated rightmost. **b,** Genome browser plots showing DNA methylation signals on one paternal GL-DMR (*H19*) and one maternal GL-DMR (*Impact*). Published coordinates and coordinates determined in our analysis are indicated in blue and gold, respectively. **c,** Heatmap showing all 859 identified DMRs. Hierarchical clustering was performed based on DNA methylation levels in oocyte and sperm from published data ^27^. **d,** Distribution of novel DMRs and known GL-DMRs over different genomic features. Gene promoters and 1,000 random sets of regions with a similar size and distribution as DMRs are shown for reference. **e,** Locus overlap analysis ^33^ of published ChIP peaks (Zfp445 and Setdb1) on known GL-DMRs and novel DMRs. **f,** Motif enrichment analysis ^38,50^ in known GL-DMRs and novel DMRs.

We next analysed available oocyte and sperm DNA methylome data ^27^ to determine the developmental origins of the 859 DMRs. Of our 778 maternal DMRs, 410 (52%) exhibited oocyte-specific or oocyte-biased DNA methylation (clusters C1 and C4, respectively) (Fig. 3c) and 62 of the 81 (76%) paternal DMRs were methylated in sperm genomes (cluster C5). Notably, 349 blastocyst-DMRs (41% of the total) were established during preimplantation development by loss of DNA methylation on one allele, typically the paternal one (cluster C2) ^1^. Thirty-seven blastocyst DMRs exhibited little or no DNA methylation in oocytes or sperm (cluster C3). These data collectively suggest that most differential DNA methylation is encoded within gamete genomes, even though DMRs may not become manifest until later in development.

Most GL-DMRs (> 75%) were located within gene bodies (Fig. 3d). For novel DMRs, we detected an overlap with gene promoters and CpG islands of ~25%. We asked which chromatin regulators might interact with the identified DMRs using two complementary bioinformatics approaches. First, we tested for overlaps between DMRs and binding regions of transcription factors and chromatin modifiers by mining 791 published ChIP-Seq datasets ^33–35^. This identified enrichments of the H3K9-specific histone-lysine methyltransferase Setdb1 and the zinc finger protein Zfp445, after subtracting overlaps generally detected in promoters (Fig. 3e). Setdb1 establishes H3K9me3, an imprint-associated chromatin mark ^36^ and Zfp445 is a primary regulator of genomic imprinting ^37^. Secondly, we searched the DMRs for matches to transcription factor DNA-binding motifs ^38^. This detected an enrichment for Zfp57, E2f5, and Nrf1 cognate sequences (Fig. 3f), which were also enriched in known GL-DMRs. Zfp57 synergistically contributes to imprint maintenance with Zpf445 ^30,37^ and Nrf1 has also been implicated in imprinted gene regulation ^39^. These findings together support the view that novel DMRs share chromatin regulatory features with known GL-DMRs.

### Associating DMRs with parent-of-origin biased expression

Integrating parent-of-origin biased blastocyst transcriptome and DNA methylome data has the potential to reveal relationships between gene expression and DNA methylation. We found that the vast majority of both nBiX and nBsX genes exhibited paternal expression with maternal 5mC at the closest DMR, similar to published imprinted genes (Fig. 4a). However, whereas substantial numbers of both published imprinted (62/134; 46%) and confirmed published imprinted (12/34; 35%) genes resided close to a DMR (< 250kb between DMR and gene), nBiX and nBsX did not show such an association (8/71 [11%] and 15/106 [14%], respectively) (Fig. 4b; Extended Data Fig. 4a and b). Moreover, even in published and confirmed published imprint gene sets, most genes (72/134 [54%] and 22/34 [65%], respectively) were not located near to a DMR, suggesting that proximal DMRs are not a defining feature of imprinted genes in blastocysts.

**Fig. 4.**
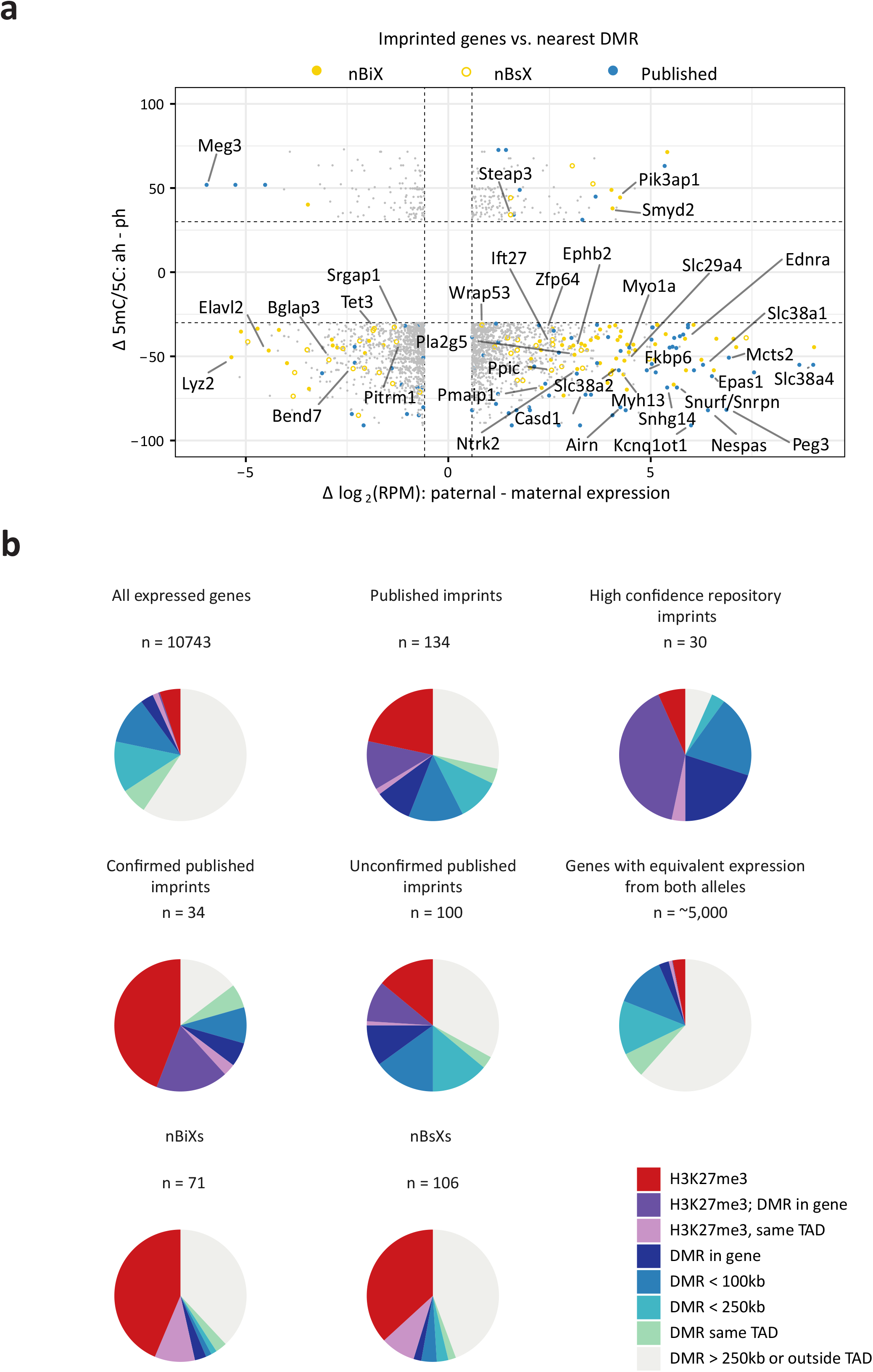
Intersecting parent-of-origin-specific DNA methylation and H3K27me3 with parental allele-specific gene expression. **a,** Plot contrasting differential DNA methylation in uniparental blastocysts (y-axis) and parent-of-origin-specific gene expression (x-axis). Published and novel imprinted genes (nBiX and nBsX) are shown in color, other genes are indicated in grey. Each dot represents one gene associated with the closest DMR. Selected genes were labelled. **b,** Pie charts showing all 10,743 robustly detected genes, published imprinted genes with expression data, published imprinted genes which were confirmed in blastocysts (intersection with BiX), published imprinted genes which are not part of the BiX, genes that are bi-allelically expressed in blastocyst, nBiX and nBsX, showing their association to different genomic features (DMRs, parent-of-origin-specific H3K27me3 on TSS). Distances of these genes from their nearest DMR are color coded. Further color codes indicate the presence of allele-specific H3K27me3 on the gene promoter (TSS +/− 5kb) or association with a DMR in the same topologically associating domain (TAD), independent of distance.

Only eight nBiX genes were within 250 kb of a DMR (Fig. 4b; Extended Data Fig. 4a). The large distance of most nBiX genes to their nearest DMR (relative to that of known imprinted genes) may be explained by long-range tertiary chromatin interactions between DMRs and nBiX loci. We addressed this possibility by utilizing HiC data from mouse ESCs ^40^ and investigated the concomitant presence of nBiX or nBsX with DMRs in the same topologically associated domain, TAD (Extended Data Fig. 4c). Even after considering DMR-gene pairs within the same TAD, only 18% of nBiX and 20% nBsX genes were potentially associated with DMRs (Fig. 4b; Extended Data Figure 4c).

Recent research suggests that parent-of-origin-specific H3K27me3 functions in specifying imprinted gene expression in preimplantation development ^9^. We therefore interrogated available datasets to map H3K27me3 to the transcription start site (TSS) of parent-of-origin specific genes ^41^. We identified parent-of-origin-specific H3K27me3 at the TSS of 741 out of all 10743 (7%) genes with expression evidence in our datasets, and 217 out of 5376 (4%) genes equivalently-expressed from both alleles, defining the genomic background. Forty-seven out of 134 (35%) published imprinted genes showed an enrichment of parent-of-origin-specific H3K27me3 on respective TSSs, which is less than the overlap with proximal DMRs (46%) (Extended Data Fig.4d). This indicates that published imprinted genes are more closely associated with DMRs than with parent-of-origin-specific H3K27me3. However, 22 out of 34 (65%) of the group of confirmed published imprinted genes exhibited parent-of-origin-specific TSS-associated H3K27me3, suggesting that H3K27me3 assists in regulating parent-of-origin-specific expression of known imprinted genes at the blastocyst stage. Both nBiX and nBsX exhibited only low levels of association to DMRs, but parent-of-origin-specific H3K27me3 peaks at promoters on par with those observed for confirmed published imprinted genes (54% and 45% for nBiX and nBsX, respectively). This suggests that there is a senior role for Polycomb, and a junior one for DNA methylation, in regulating (n)BiX gene expression. Interestingly, we found that 5 out of 34 confirmed published imprinted genes (15%), and 27 out of 71 nBiX genes (38%) were neither associated with parent-of-origin-specific H3K27me3 nor with a proximal DMR.

### Functional dependence of novel and known imprinted gene expression on maternal H3K27me3 and DNA methylation

Association with an epigenetic mark does not necessarily signify causality for mediating imprinted expression. To address whether the relationship was causal, we took advantage of available datasets mapping allele-specific expression of mouse morulae carrying a maternal deletion of either *Dnmt3l* (*mDnmt3l* matKO) or *Eed* (*mEed* matKO) ^42,43^. We reasoned that genes showing allelically skewed expression in control embryos, which is lost or reduced upon maternal *Dnmt3l* (*mDnmt3l*) or *Eed* (*mEed*) depletion, would be regulated by DNA methylation or H3K27me3, respectively. Accordingly, we compared allele-specific expression in our datasets with the response of allelic skewing upon *mDnmt3l* or *mEed* depletion in morula datasets (Fig. 5a-c). Whereas confirmed published imprinted genes showed only limited response to loss of *mDnmt3l*, there was a strong reduction of parent-of-origin skewing in *mEed* KO morulae (Fig. 5a). For published imprinted genes that did not show parent-of-origin-specific expression in our datasets, there was no response to loss of either *mDnmt3l* or *mEed* (Fig. 4b). However, nBiX genes exhibited behaviours similar to those of confirmed published imprinted genes (Fig. 5c). We then mapped physical associations with DMRs (within 250 kb or in the same TAD) and parent-of-origin-specific H3K27me3 onto the KO expression data (Fig. 5a-c). This revealed an enrichment for parent-of-origin-specific H3K27me3-decorated TSSs among genes responsive to loss of *mEed* among the groups of published confirmed imprinted and nBiX genes, but not for unconfirmed published imprinted genes. Together, these findings suggest a major role for H3K27me3 in establishing and/or maintaining parent-of-origin specific expression in pre-implantation development.

**Fig. 5.**
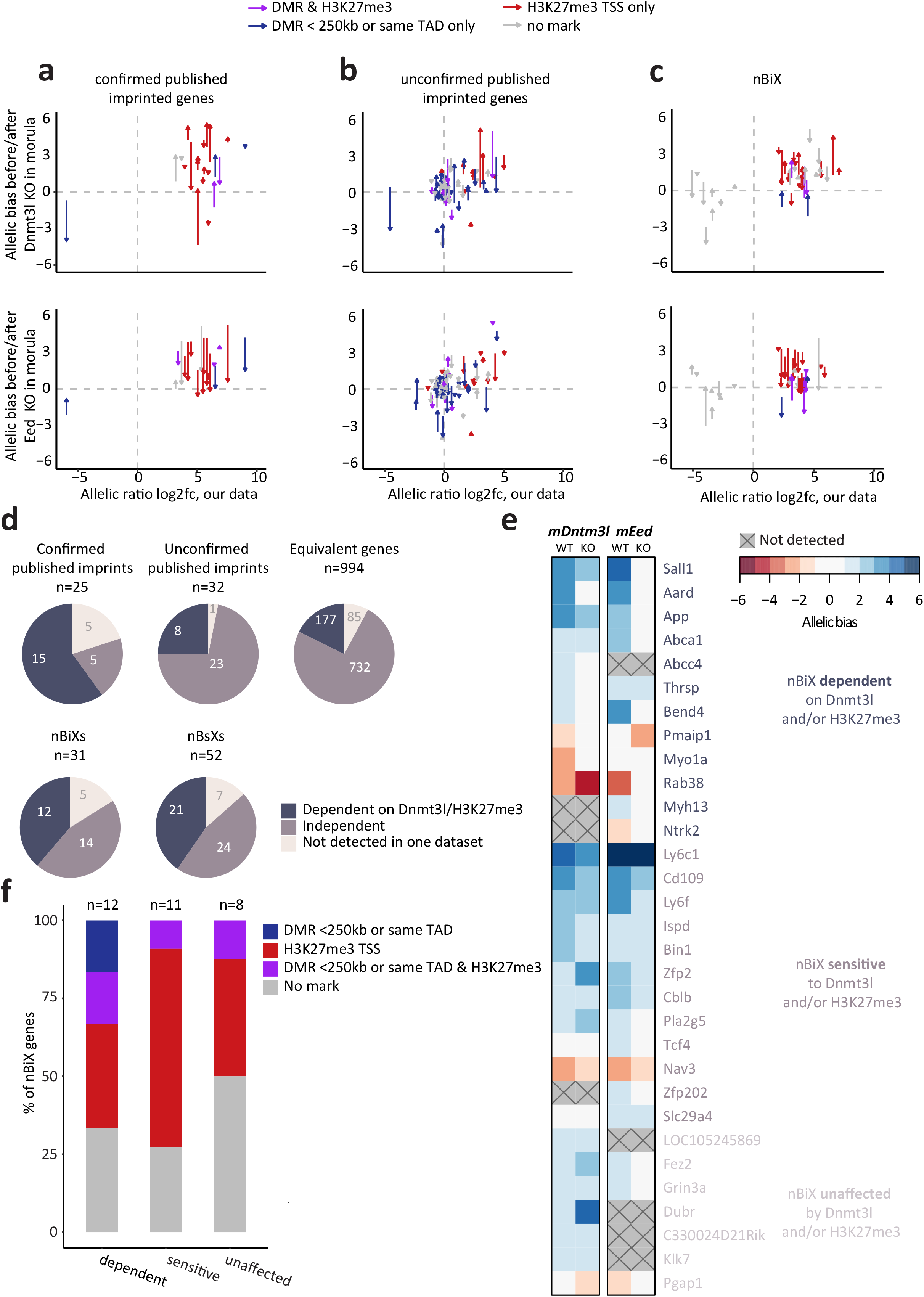
Functional dependence of novel candidate genes on maternal H3K27me3 or maternal DNA methylation. **a-c,** Differences in allele-specific expression induced by maternal knockout (mKO) of *Dnmt3l* (top) and *Eed* (bottom) ^42,43^. Confirmed published (a), unconfirmed published (b), and novel imprinted genes (nBiX; c) are shown in separate panels. In the plots, each arrow points from the allele-specific expression ratio (ratio of SNP-containing maternal and paternal reads; log2 fold change) in wild-type toward the expression ratio of the same gene in mKO morulae. **d,** Pie charts showing how many genes within the represented group (nBiX, nBsX, confirmed published imprints, unconfirmed published imprints, and equivalent genes) lose parent-of-origin specific expression following maternal deletion of either *Dntm3l* or *Eed* in morulae (“dependent” on Dnmt3l / H3K27me3; allelic bias > |1| before and < |1| after maternal depletion of epigenetic regulator with a log2fc > |1| between wt and mutant.). Colour indicates genes that were dependent (blue) or independent (grey) of *Dnmt3l* and *Eed*. Genes that were only detected in one dataset and did not show dependence are shown in light grey. **e,** Heatmap showing the allelic bias for the indicated nBiX genes in WT or mKO morulae. **f,** Bar chart showing the relative percentages of genes associated with DMRs (within 250 kb or in the same TAD), genes with allele-specific H3K27me3 mark near the transcription start site (TSS +/− 5kb), genes associated to both marks and gene associated to neither of them for the indicated groups of genes (nBiX dependent on Dnmt3l and/or H3K27me3, nBiX sensitive to Dnmt3l and/or H3K27me3 and nBiX apparently unaffected by either mechanism).

To identify those imprinted genes that show dependence on *mEed* or *mDnmt3l*, we considered that for dependent genes, parentally biased expression in wild-type (wt) morulae would be lost in mutant morulae (Extended Data Figure 5a, b). This analysis showed that imprinting was dependent on either *mEed* or *mDnmt3l* for 60% of published confirmed imprinted genes (Fig. 5d; Extended Data Fig. 5b). Expression of 40% of nBiX and nBsX genes were dependent on maternal H3K27me3 or maternal DMRs. However, expression of some 50% of novel imprinted genes were apparently H3K27me3- and DMR-independent.

Functional contribution of *Dnmt3l* or *Eed* to parent-of-origin-specific expression in blastocysts is not necessarily predicted to result in complete loss of allele-specific expression in *mEed* or *mDnmt3l* mutant morulae, but may merely cause a reduction in allelic bias. This might, for example, be the case for genes that are coregulated by multiple chromatin-regulatory mechanisms. We therefore asked whether expression of genes that fell below our strict cut-off exhibited loss of either known imprint-defining epigenetic mark. We defined genes as ‘sensitive’ if they exhibited allelic skewing in wt morulae and reduced allelic bias upon loss of mEed or mDnmt3l in morulae, even if some parental bias could still be detected. Eleven out of 19 nBiX genes and 16 out of 31 nBsX genes not showing dependence were sensitive to loss of either Dnmt3l or Eed (Fig. 5e; Extended Data Fig. 5c). The majority of strictly Eed*/*Dnmt3l-dependent, and -sensitive nBiX genes were marked by DMRs or allele-specific H3K27me3 (Fig. 5f). Only for four nBiX and six nBsX genes we could detect neither Eed/Dnmt3l dependence or sensitivity, nor the physical presence of known imprint-associated chromatin modifications. In summary, this shows that most novel blastocyst imprinted genes are regulated by parent-of-origin specific DNA methylation or H3K27me3. The data suggest that the Polycomb machinery, both at levels of gene expression and chromatin, plays a major role in establishing and/or maintaining imprinted expression of known and novel imprinted genes in the blastocyst.

### Identification of imprinted gene clusters in blastocysts

Imprinted genes are known to reside in genomic clusters regulated by *cis*-acting imprinting control regions (ICRs) ^44–46^. We therefore searched for clusters containing at least two published or novel (nBiX or nBsX) imprinted genes within 250 kb, yielding 32 potential ICR clusters (Fig. 6a). Twelve clusters contained at least one published imprinted gene that exhibited uniparental expression in blastocysts (Fig. 6a, clusters #1-12). Eight of these also contained a DMR within at least one of its associated genes or less than 10 kb away. One additional cluster encompassing the *Slc38a1* gene contained a DMR within the same TAD. Strikingly, ten clusters of published imprinted genes (including the *Igf2* cluster) lacked significant expression bias in blastocysts, despite exhibiting a DMR within the gene body in nine of these ten cases (Fig. 6a, clusters #13-22). This suggests that in some cases, differential DNA methylation and parent-of-origin-specific expression is unlinked. A subset of six published imprinted genes in these clusters (including *Commd1* and *Grb10*) contained both a DMR and parental-allele-specific H3K27me3 on their TSSs, but apparently neither of these epigenetic marks elicited allele specific gene expression. Our analyses extended five known imprinting clusters by identifying novel imprinted genes close to published examples (Fig. 6a, cluster #23-27). Although these clusters were devoid of proximal DMRs, they were all associated with parent-of-origin-specific H3K27me3. In addition, five clusters were identified comprising exclusively nBiX genes. Four contained at least two protein-coding genes (Fig. 6a, cluster #28-31), and one contained a protein coding gene and a ncRNA (Fig. 6a, cluster #32). Neither of these clusters possessed a DMR within 250kb or in the same TAD, but two clusters were associated with allele-specific H3K27me3 on the TSS of at least one cluster member. Three nBiX clusters were not associated with either DMRs or maternal H3K27me3.

**Fig. 6.**
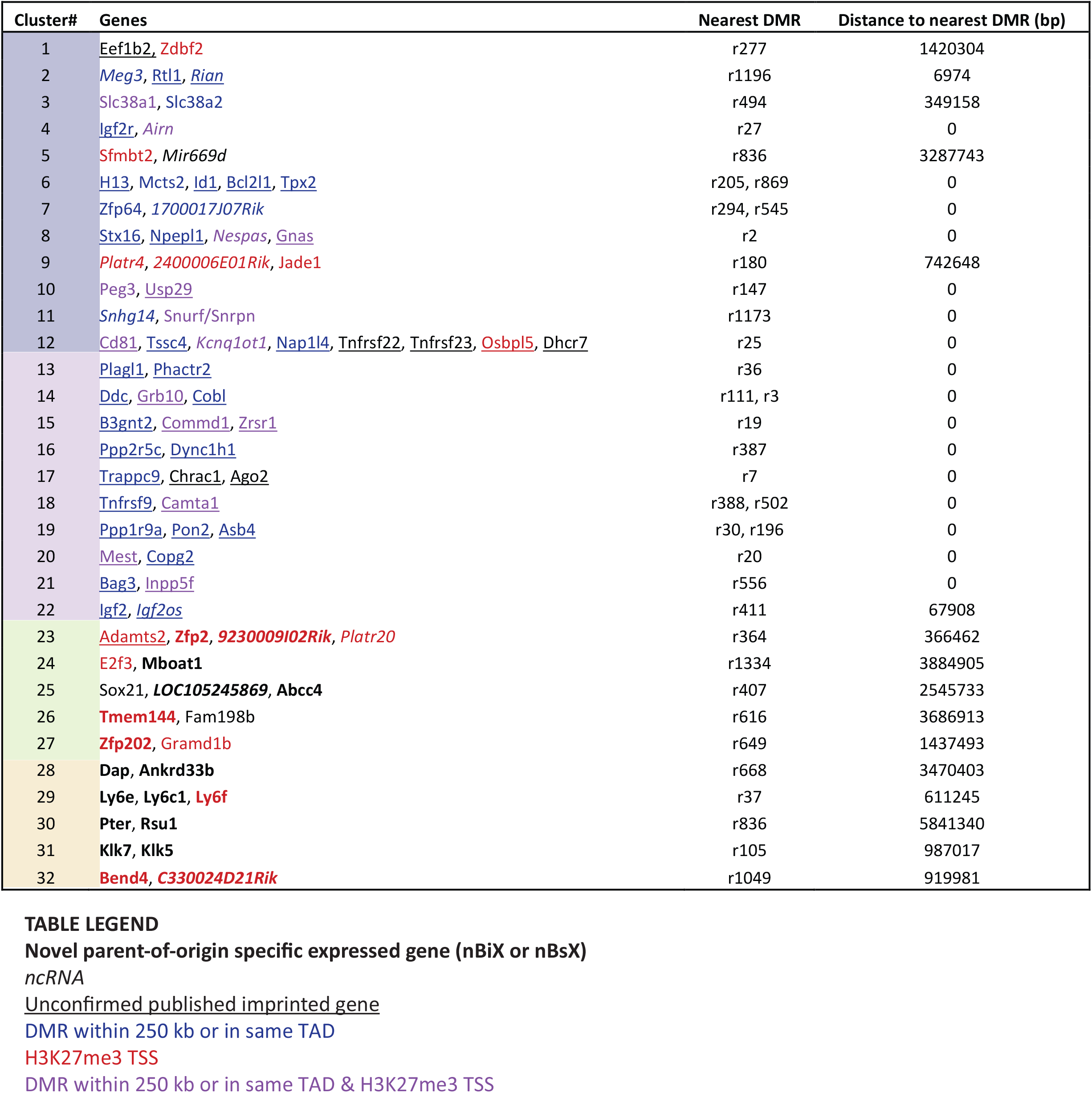
Novel imprinting clusters and novel genes in known clusters. **a,** Gene clusters (distance between features in cluster ≤ 250kb) consisting of published imprinted genes with one or more confirmed BiX per cluster (#1-12, blue background); clusters of published imprinted genes without confirmed parent-of-origin-specific expression (#13-22, violet background); clusters consisting of published imprinted genes with at least one novel imprinted gene (#23-27, green background) and clusters containing only novel candidate genes (#28-32, yellow background). Colour code: black: published imprinted genes with parent-of-origin specific expression evidence in blastocyst (published BiX or BsX); underlined, published imprinted gene without evidence for parent-of-origin specific expression in blastocyst; bold, novel imprinting candidate (nBiX or nBsX); blue, gene associated with a proximal DMR (within 250 kb or in the same TAD); red, gene with allele-specific H3K27me3 mark on its TSS; italic, ncRNA. Distances to the nearest DMR (and identifiers) are indicated.

## DISCUSSION

By combining allele-specific transcriptomics and uniparental DNA methylome profiling, we have delineated the imprinting status of blastocyst-stage mouse embryos, identifying 859 parent-of-origin-specific DMRs and 106 genes with parent-of-origin-specific allelic bias (nBsX genes), and 71 novel imprinted genes with parent-of-origin-specific expression with an allelic ratio of 70:30 or greater (nBiX genes). Parental expression bias was evident in blastocysts for only 34 of 134 published imprinted genes and we detected statistically indistinguishable expression of both alleles for 24 published imprinted genes. Of those, four including *Commd1* exhibited significant biallelic expression in blastocysts despite each having a DMR within their gene-bodies, showing that differential DNA methylation is not sufficient to guarantee uniparental expression. This suggests that hitherto unappreciated tissue- and stage-specified programmes underlie the regulation of imprinted gene expression.

Published imprinted genes represented in the BiX dataset exhibited different DNA-methylation profiles compared to published imprinted genes not showing significant parent-of-origin-specific expression (Fig. 4b). This points to the existence of at least two broad classes of known imprinted genes: i) those with closely-associated DMRs, which are likely to be uniparentally expressed in blastocysts, and ii) those with more distal DMRs, which are less likely to exhibit parent-of-origin specific expression in blastocysts.

Many DNA methylation-based imprints are GL-DMRs, and most (55%) of the DMRs we detected in blastocysts had indeed also been identified in gamete-specific DNA-methylation analysis. A further 41% of blastocyst DMRs were generated by parent-of-origin-specific loss of DNA methylation on one (mainly the paternal) parental allele. We also detected a gain of allele-specific DNA methylation at some loci that were unmethylated in gametes, suggesting that parent-of-origin specific DNA methylation in blastocysts can be encoded in gametes independently of DNA methylation and later decoded to allow DNA methylation during preimplantation development.

Despite reported imprint instability in 2i culture ^47^, diploid ESCs relatively robustly sustained DMRs with the exceptions of the *Gnas* and *Liz1*/*Zdbf2* loci, at least within the passaging range (8 to 20 passages) used in our assays. We also observed a tendency for haploid, but not diploid ESCs to lose DMRs, in contrast to stable imprint maintenance in human haploid parthenogenetic ESCs ^48^. Whether this difference in imprint stability in haploid ESCs reflects species, culture or cell state (*e.g.* naïve *vs* primed pluripotent) differences is unclear.

Our data also show that H3K27me3- and DMR-based imprinting mechanisms regulate overlapping but largely distinct gene sets. Indeed, some well-studied imprinted genes, including *Airn*, *Snrf*/*Snrpn*, *Peg3*, *Fkbp6*, *Nespas*, *Kcnq1ot1* and *Slc38a1* harbour both proximal DMRs and H3K27me3 peaks on their TSSs. However, imprinted expression of these genes is not necessarily directly dependent on H3K27me3; it depends on maternal *Dnmt3l* for *Peg3*, *Nespas*, *Fkbp6* and *Kcnq1ot1*. This suggests that overlapping chromatin profiles of allele-differential DNA methylation and H3K27me3 do not always translate into functional redundancy and that additional layers of regulation exist that dictate dependence of imprinted expression on H3K27me3 or DMRs.

Clustering of imprinted genes facilitates coordinated parent-of-origin-specific expression control, such that a given ICR can regulate the expression of multiple genes. We identified five novel imprinted gene clusters and new members of multiple known clusters. All novel (and all expanded) clusters lacked blastocyst DMRs detected within 250kb or present in the same TAD. Six out of the 10 ‘non-DMR clusters’ contained at least one gene associated with H3K27me3, suggesting that their imprinting is controlled *via* allele-specific PRC2 mediated histone modification at E4.5.

In sum, this work provides a detailed compendium that identifies novel imprinted genes and imprinting clusters. It reveals a major contribution of Polycomb mediated imprint control in blastocysts (Fig. 7), suggesting that imprint regulation in preimplantation embryos is achieved by both H3K27me3- and DMR-dependent mechanisms. The implication is therefore that there exist different tiers of mechanistically distinguishable, potentially stage-specific imprinting that must be integrated for the healthy development of preimplantation embryos and beyond.

**Fig. 7.**
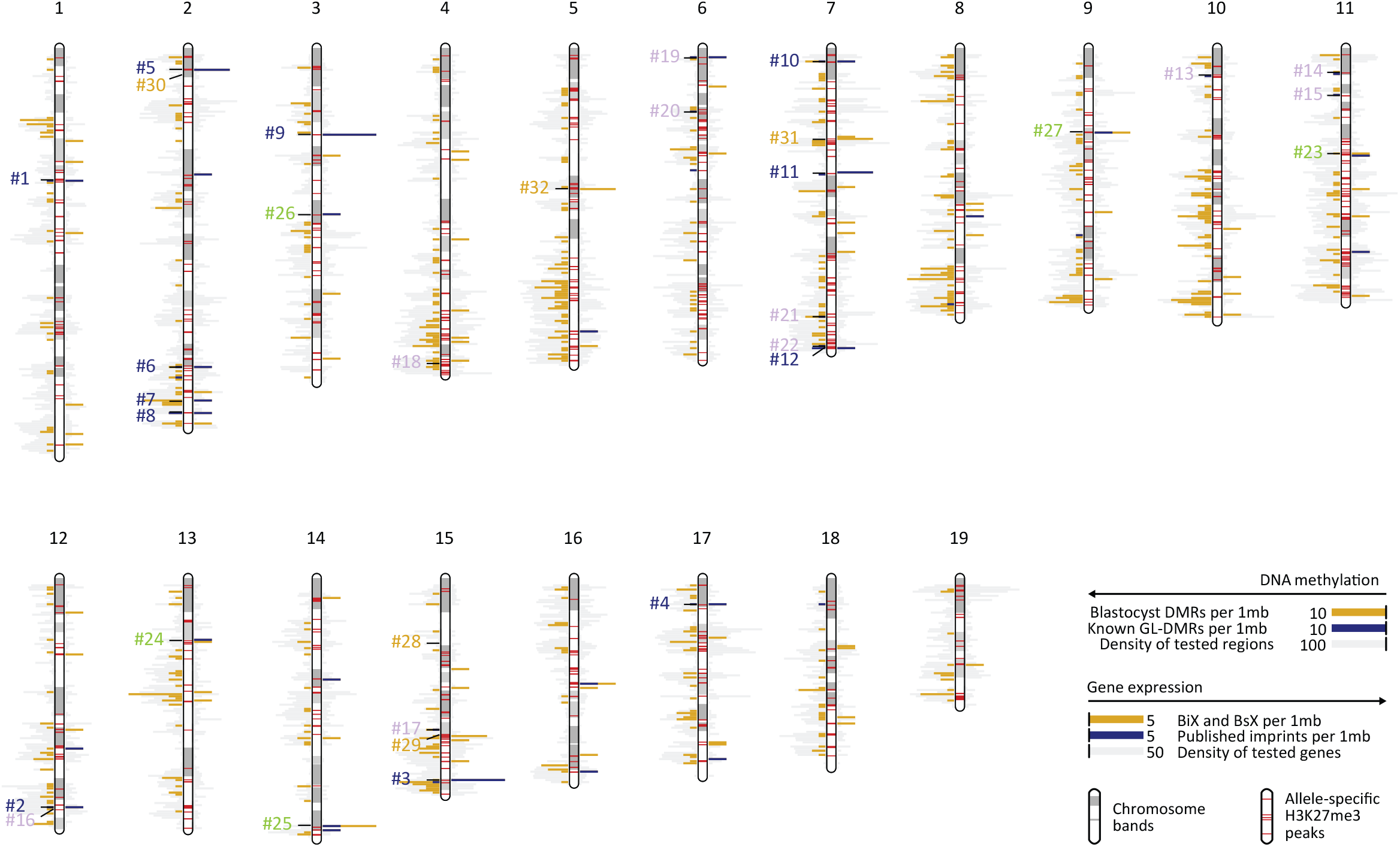
Genome-wide overview of BIX and DMRs. Visualization of the chromosomal locations of imprinted genes and chromatin marks. Novel blastocyst-specific DMRs are plotted as bars to the left in gold, known GL-DMRs are shown in blue. The density of tested regions (regions with reads in μWGBS) are plotted in grey. Parent-of-origin expression bias is shown on the right. nBiX and nBsX are plotted in gold, known imprinted genes in blue. The density of all robustly expressed genes are plotted in grey. All clusters of Figure 6 are indicated. The locations of all allele-specific H3K27me3 peaks are shown as red bands overlaid on the chromosome ideograms.

## AUTHOR CONTRIBUTIONS

LS performed experiments and analysed data. FH performed biocomputational experiments and analysed data. FTT performed allele specific RNAseq analysis with support from FP and SH. AB supervised RNAseq analysis. TS generated androgenetic and parthenogenetic embryos and performed ‘semicloning’ experiments. MA performed embryo immunofluorescence assays. MF and CB performed and supervised SMARTseq2 and μWGBS analysis. JR provided experimental support to LS. XM performed TAD analysis supervised by EDH. ACFP and ML conceived and supervised the study, provided funding an analysed data. ACFP and ML wrote the paper together with LS, FH and FTT with input from all authors.

**Extended Data Fig. 1.**
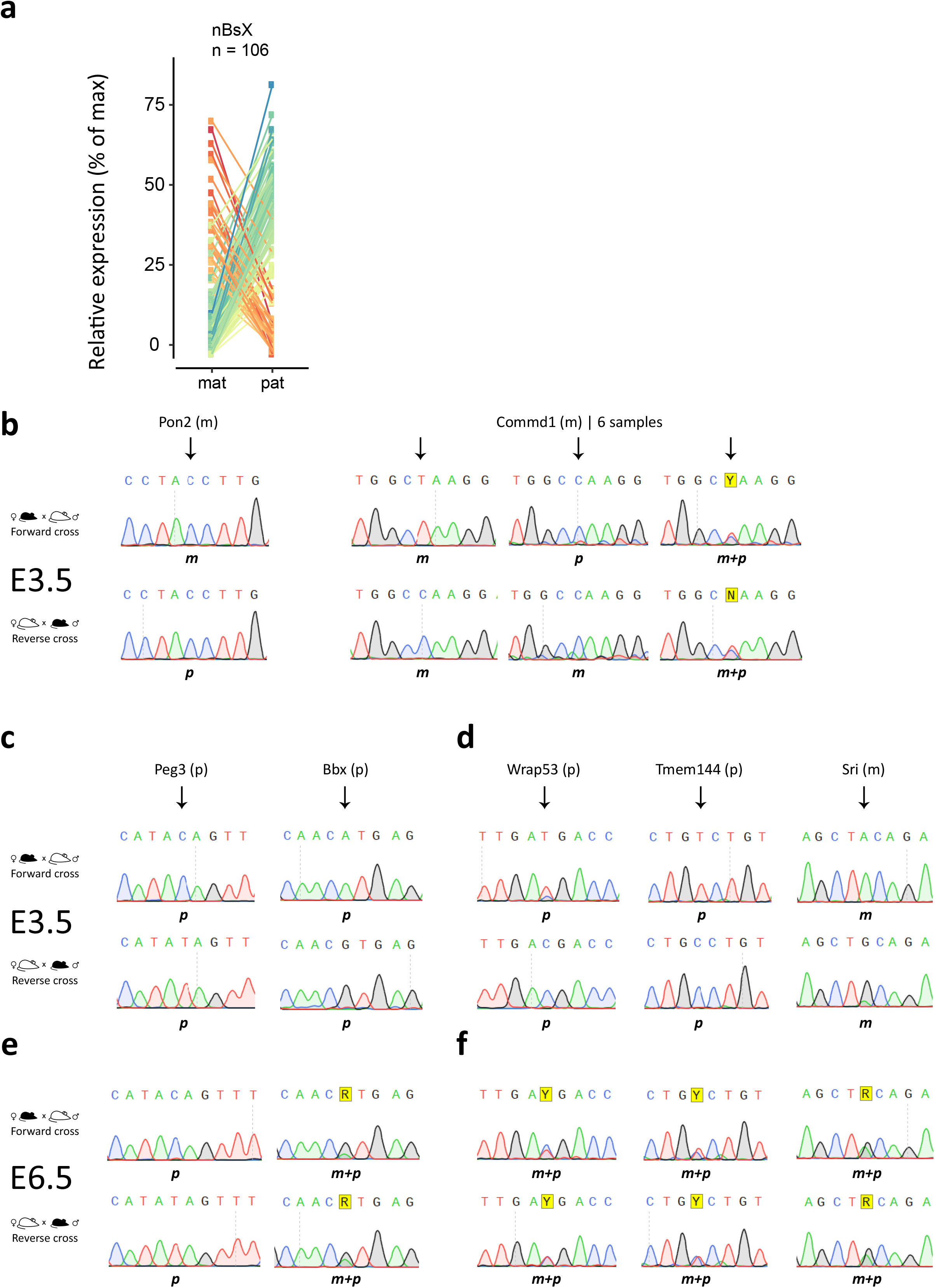
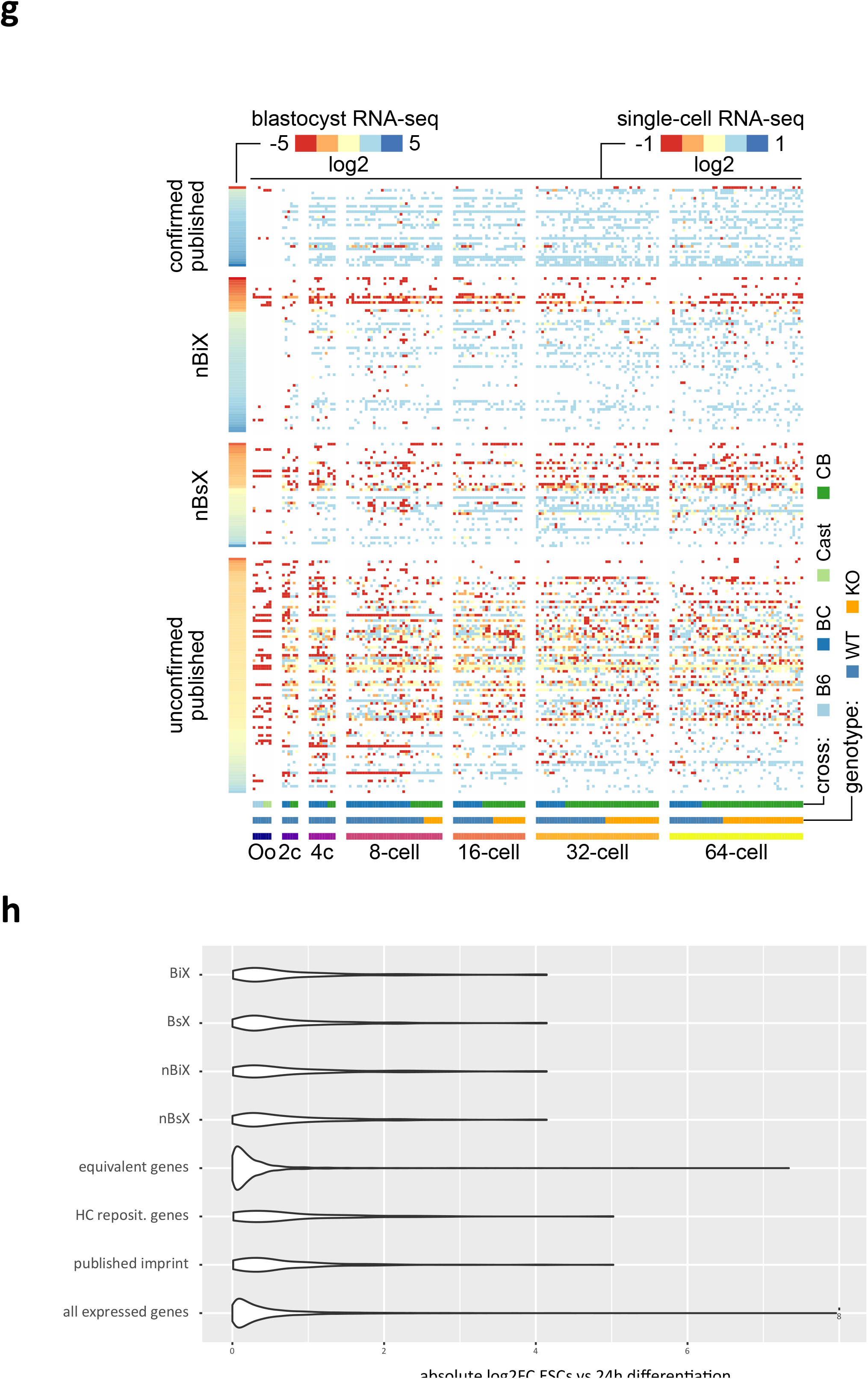
Parent-of-origin-specific gene expression analysis in blastocysts (related to Fig. 1) **a,** Distribution of SNP-containing RNA-seq reads between maternal and paternal alleles in nBsX genes. **b,** RT-PCR and Sanger sequencing-based analysis of allele-specific expression of *Pon2* and *Commd1* using E3.5 embryo samples; published imprinted genes for which we could not confirm parent-of-origin-specific expression in blastocysts. **c,** RT-PCR and Sanger sequencing-based validation of allele-specific expression of the confirmed published imprinted genes *Peg3* and *Bbx* at E3.5 **d,** RT-PCR and Sanger-sequencing-based validation of nBsX genes in E3.5 embryos. **e,** RT-PCR and Sanger sequencing-based validation of allele-specific expression of the confirmed published imprinted genes *Peg3* and *Bbx* at E6.5 **f,** RT-PCR and Sanger-sequencing-based validation of nBsX genes in E6.5 embryos. **g,** Heatmap visualizing single cell sequencing data from Borensztein et al., 2017 showing parent-of-origin specific expression at stages during preimplantation development. **h,** Violin plot showing absolute log2fold changes of indicated groups between ESCs (2i) and cells at an early stage of differentiation (24h after 2i withdrawal) ^24^.

**Extended Data Fig. 2.**
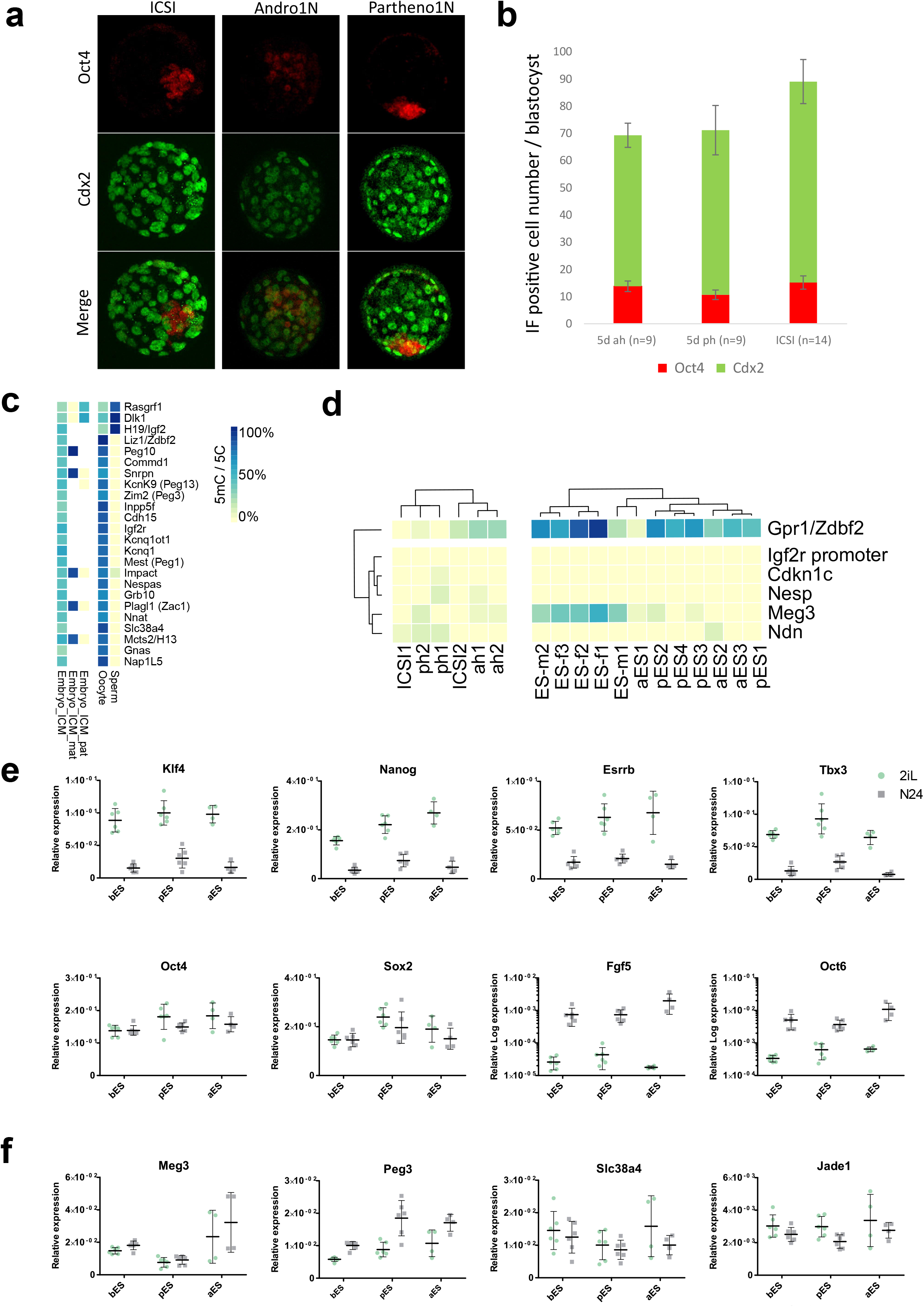
DNA methylation analysis confirms genomic imprints at known GL-DMRs (related to Fig. 2) **a,** Immunofluorescence analysis showing Oct4 and Cdx2 expression in ICSI, androgenetic (Andro1N) and parthenogenetic (Partheno1N) blastocysts. **b,** Quantification of data shown in (a). Errors bars show standard deviation between samples. Sample numbers are indicated. **c,** Heatmap showing DNA methylation signal over 24 known GL-DMRs in previous data ^27^. **d,** Heatmap showing DNA methylation levels over known somatic DMRs in indicated samples. **e,** RT-qPCR analysis showing expression levels of pluripotency and early differentiation markers in two androgenetic, three parthenogenetic, and three biparental ESCs cultured in 2i (left in each pair), and 24h after 2i withdrawal induced release into differentiation. Mean and standard deviation values for 2 independent experiments are shown. **f,** RT-qPCR analysis showing expression levels of four published imprinted genes in androgenetic, parthenogenetic and biparental ESCs cultured in 2i and 24h after 2i withdrawal as in(e).

**Extended Data Fig. 3.**
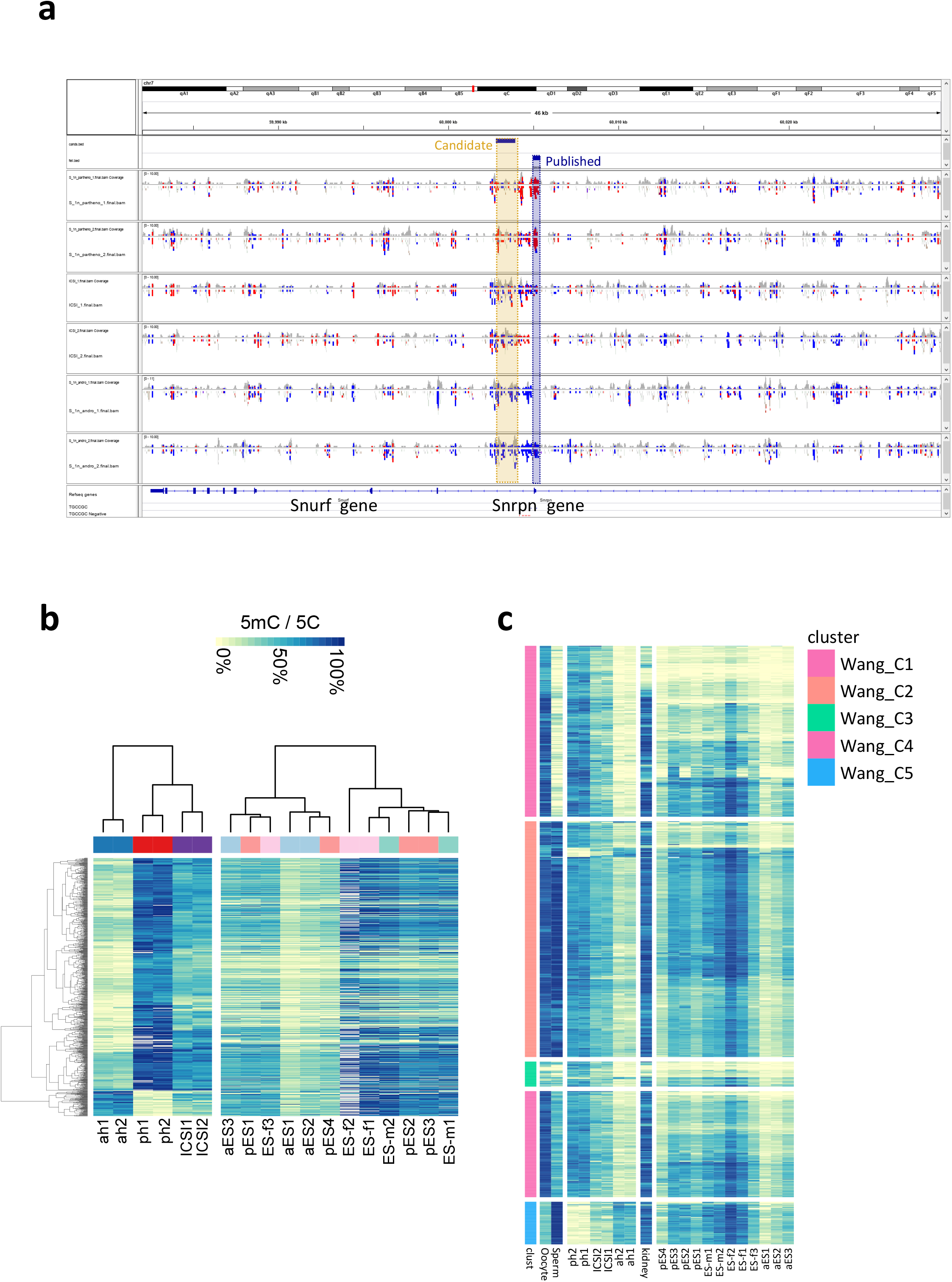
Novel DMRs in uniparental embryos (related to Fig. 3) **a,** Genome browser plot showing DNA methylation signals at the *Snrp*/*Snrf* locus. Published DMR coordinates and coordinates determined in our analysis are indicated in blue and gold, respectively. **b,** Heatmap showing clustering of all 859 DMRs in embryos (left) and ESC lines of various parental provenance. Clustering of DMRs was performed on blastocyst data only. **c,** Heatmap showing all 859 identified DMRs. Clustering was performed based on DNA methylation levels in oocyte and sperm from published datasets. ESC and somatic cell DNA methylation data extend analysis from Figure 2c.

**Extended Data Fig. 4.**
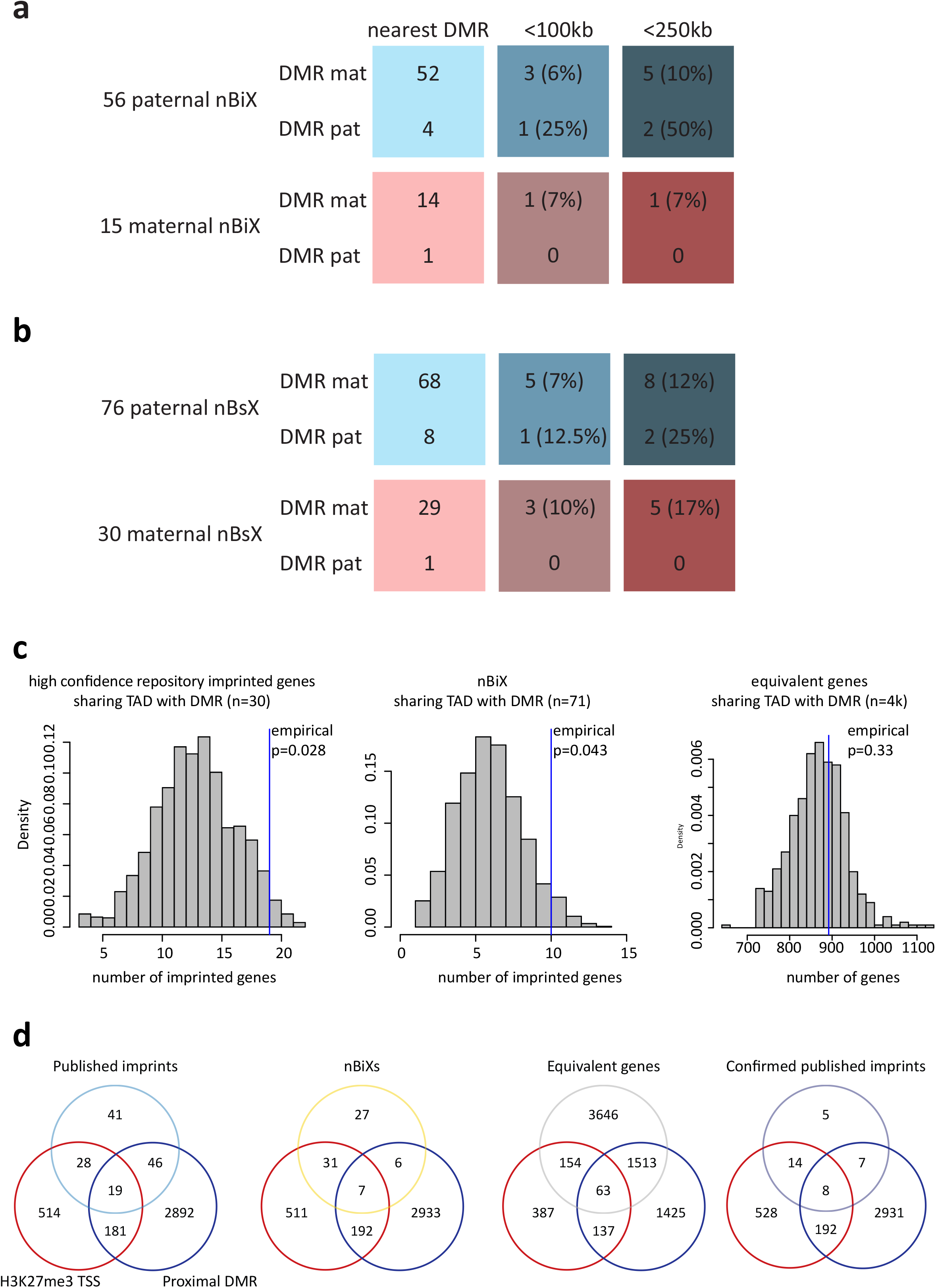
Integrated expression and methylome analysis (related to Fig. 4) **a-b,** Charts showing association between 71 nBiX genes or 106 nBsX and DMRs. Paternal nBiX/nBsX genes are shown on blue background. Increasing color intensity indicates decreasing distances of respective gene group from nearest DMRs. **c,** In-silico-derived distribution of DMR-gene pairs within the same TAD for high confidence repository imprinted genes, nBiX genes and equivalent genes. Observed value for each gene group is indicated by a blue line. Empirical p-values are indicated. **d,** Venn diagrams showing the overlaps with proximal DMR (within 250 kb or in the same TAD) and H3K27me3 on gene TSS for all detected published imprinted genes, published imprinted genes with allele-specific expression in blastocyst, nBiX genes and genes bi-allelically expressed in blastocyst.

**Extended Data Fig.5.**
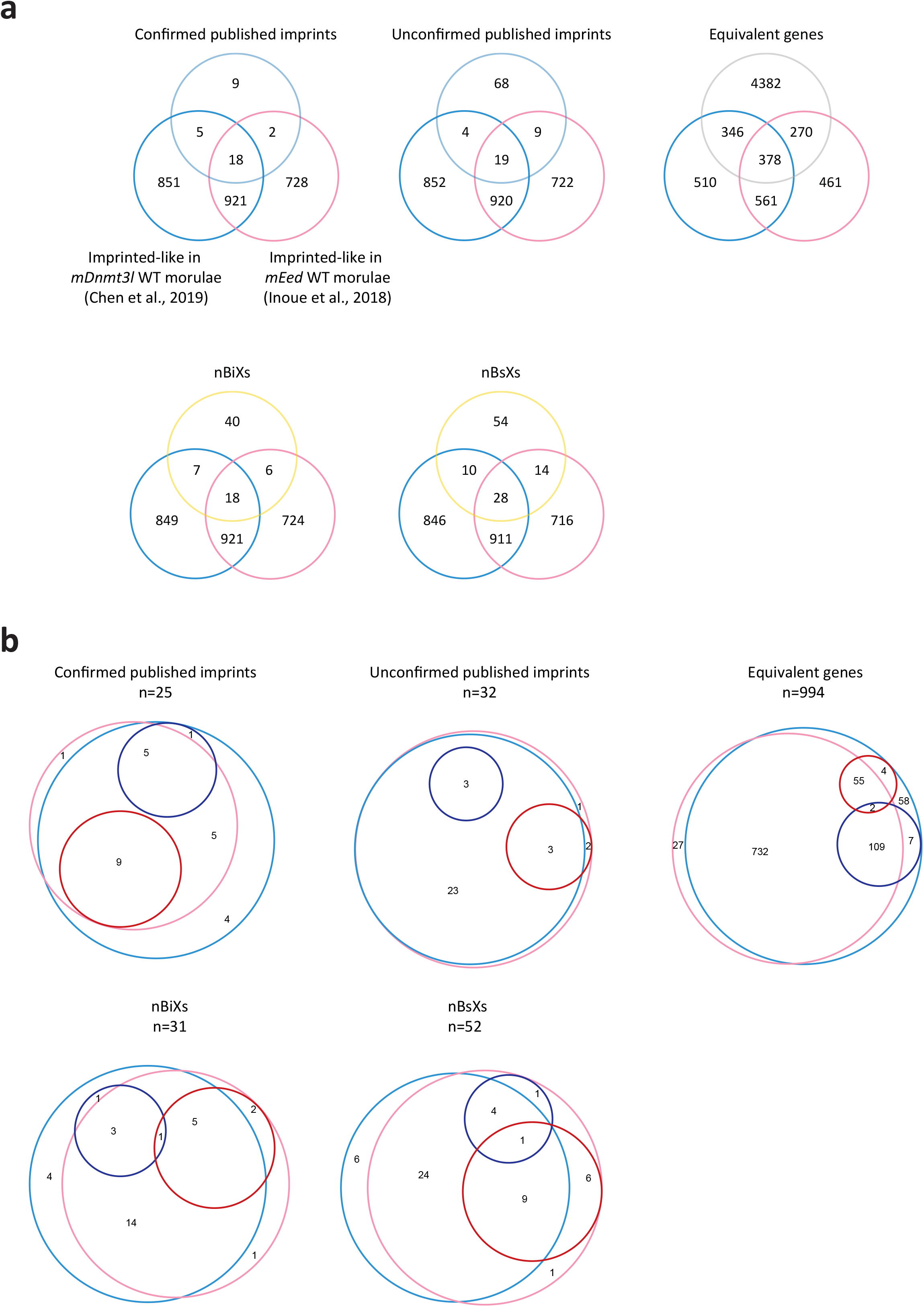
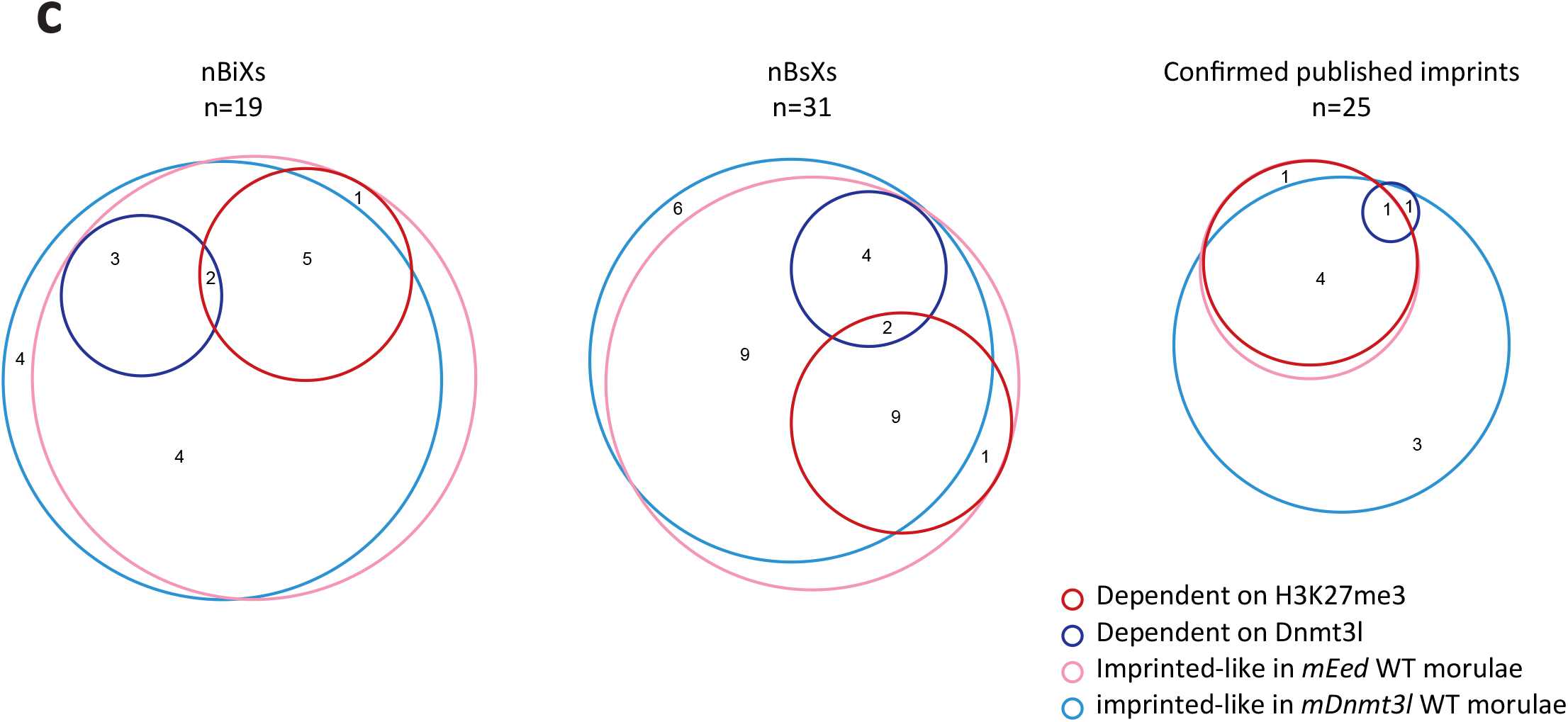
Functional dependence of novel candidate genes on maternal H3K27me3 or maternal DNA methylation (related to Figure 5) **a,** Venn diagrams showing overlap between the indicated groups of genes (nBiX, nBsX, confirmed published imprinted genes, unconfirmed published imprinted genes, and equivalent genes) and genes with imprinted-like expression in control morulae from Chen et al., 2019 (light blue) or from Inoue et al., 2018 (pink circle). **b,** Venn diagrams showing overlap between genes with imprinted-like expression in control morulae from Chen et al., 2019 (light blue) and from Inoue et al., 2018 (pink), and genes that loose the imprinted-like status upon maternal deletion of *Dnmt3l* (genes dependent on Dnmt3l, blue) or *Eed* (genes dependent on H3K27me3, red) for the indicated groups of genes (nBiX, nBsX, confirmed published imprinted genes, unconfirmed published imprinted genes, and equivalent genes). **c,** Venn diagrams showing overlap between genes with imprinted-like expression in control morulae from Chen et al., 2019 (light blue) and from Inoue et al., 2018 (pink circle) that showed no dependence to H3K27me3/Dnmt3l as defined for (**b**), and genes with reduced imprinted-like status upon maternal deletion of Dnmt3l (genes sensitive to Dnmt3l, blue circle) or Eed (genes sensitive to H3K27me3, red circle) for the indicated groups of genes (nBiX, nBsX and confirmed published imprinted genes).

## METHODS

### Animals

Animal procedures complied with the statutes of the Animals (Scientific Procedures) Act, 1986, approved by the University of Bath Animal Welfare and Ethical Review Body and the Biosciences Services Unit. Wild-type mouse strains were bred from stocks in-house or otherwise supplied by Charles River (L’Arbresle, France) or MRC Harwell. The strain B6D2F1 (C57BL/6xDBA2) was generally used as a source of unfertilized metaphase II (mII) oocytes. Some parthenogenotes were produced from *Gt(ROSA)26Sor*^*tm4(ACTB-tdTomato,-EGFP)Luo*^ (mT) oocytes and we generated a 129/Sv-J line carrying a single, ubiquitously-expressed *pCAG-eGFP* transgene (129/Sv-J-eGFP^+/−^; Suzuki *et al*., 2014) and used sperm from hemizygotes to generate androgenetic haploid embryos for ES cell derivation. Recipient surrogate mothers were of the strain ICR (CD-1) in embryo transfer.

### Oocytes

Oocyte collection was essentially performed as described previously ^51,52^. Briefly, 8-12-week-old B6D2F1 females were superovulated by standard sequential injection with 5 IU of pregnant mare serum gonadotropin (PMSG) and 5 IU human chorionic gonadotropin (hCG) ~48 h apart. Oocyte-cumulus complexes were collected into M2 medium (Specialty Media, USA) and dispersed with hyaluronidase to denude mII oocytes, which were washed and cultured in kalium simplex optimized medium (KSOM; Specialty Media, USA) equilibrated in an incubator at 37°C containing 5% (v/v) humidified CO_2_ in air until required.

### Sperm

Preparation of sperm from 129/Sv-J-eGFP^+/−^ males was essentially as described previously (Suzuki *et al*., 2014). Epidydimides from ~12-week-old males were minced with fine scissors in nuclear isolation medium (NIM; 125 mM KCl, 2.6 mM NaCl, 7.8 mM Na_2_HPO_4_, 1.4 mM KH_2_PO_4_, 3.0 mM EDTA; pH 7.0) and sperm allowed to disperse. The sperm were washed in NIM and treated in NIM containing 1.0% (w/v) 3-[(3-cholamidopropyl) dimethylammonio]-1-propanesulfonate (CHAPS) at room temperature. The suspension was gently pelleted and sperm resuspended in ice-cold NIM and held on ice until required. Just prior to ICSI, 50 μl of the sperm suspension was mixed with 20 μl of a solution of 12% (w/v) polyvinylpyrrolidone (PVP, average *M*_r_ ≈ 360,000; Sigma, UK).

### Production of uniparental haploid embryos

To establish androgenic haploid ES (ahES) cell lines, sperm from 129/Sv-J-eGFP^+/−^ hemizygous males were injected using a piezo-actuated micromanipulator (Prime Tech Ltd., Japan) into B6D2F1 mII oocytes enucleated as described previously (Wakayama *et al*., 1998): mII oocytes were placed in M2 medium containing 5 μg/ml cytochalasin B and spindles were removed. At least 1 h post-enucleation, sperm heads were injected followed by culture in KSOM for 6 h (37°C, humidified 5% CO_2_ [v/v] in air) before recording the morphology of the resultant embryos. Embryos were separated according to whether they possessed a single second polar body (Pb_2_) and pronucleus (pn) and culture was continued for 3-4 days to be utilised for ahES cell derivation. Parthenogenetic embryos were derived by strontium chloride triggered oocyte activation in calcium free medium followed by *in vitro* culture to the blastocyst stage in KSOM.

### Parthenogenesis

Activation of membrane Tomato homozygous (*mT*^+/+^) transgenic or 129/SvJ oocytes to produce parthenogenetic haploid embryos was by exposure to medium containing 10 mM SrCl_2_, 16-17.5 h post-hCG, essentially as described ^53^.

### Sperm microinjection (ICSI)

When required, ~50 μl of sperm suspension was mixed with 20 μl of polyvinylpyrrolidone (PVP, average *M*_r_ ≈ 360,000; Sigma-Aldrich) solution (15% [w/v]) and sperm injected (ICSI) into oocytes in a droplet of M2 medium, within ~60 min, essentially as described ^51^. Injected oocytes were transferred to KSOM under mineral oil equilibrated in humidified 5% CO_2_ (v/v air) at 37°C for embryo culture.

### Establishment and culture of androgenetic and parthenogenetic haploid ES cells

Haploid ES cells were established and cultured in 2i/LIF medium as previously described ^13,14^. Androgenetic haploid ES cells be maintained only in the presence of LIF and not in 2i alone. Both androgenetic and parthenogenetic haploid ES cells were recurrently sorted based on DNA content, either by Hoechst staining (15μg/ml for 15 min @ 37°C) or based on FCS/SSC parameters ^54^. The following ESC lines were used in this study: ES-f1 at p 20 (Rex1::GFPd2 reporter cell line (Leeb 2014), ES-f2 at p17 (129/B6 F1 hybrid female line), ES-m1 at p16 (E14TG2a male ESC line, ES-m2 p8 (male 129 derived ESC line), ES-m3 p8 (male ES cell line of mixed background carrying a floxed but intact Mek1 allele), pES1 p12 (haploid parthenogenetic Rex1::GFP reporter cell line, 129 background ^55^, pES2 p8 (haploid parthenogenetic ESC line P1 from a 129 background), pES3 p8 (haploid parthenogenetic ESC line T8, carrying a constitutive dtTomato reporter from a 129 background), pES4 p12 (haploid parthenogenetic ESC line H129-1 from a 129 background ^14^), aES1 p12 (haploid androgenetic ESC line A6GFP from a 129 background, carrying a constitutively active GFP transgene), aES2 p8 (haploid androgenetic ESC line A7 from a 129 background), aES3 p8 (haploid androgenetic ESC line A11 from a 129 background).

### Androgenetic haploid ES cell nucleus injection

Suspensions ahES were mixed with 20 μl of a solution of 12% (w/v) PVP and injected into mII oocytes essentially as described previously (Wakayama et al., 1998). Following a recovery period of 10-15 min, injected oocytes were activated by incubation at 37°C under 5% (v/v) humidified CO_2_ in air for 2-4 h in calcium-free CZB-G medium supplemented with 10 mM SrCl_2_ (REF). After 6~8 h, the number of Pb_2_ and pn in embryos was determined and those with a single Pb_2_ and two pn (Pb_2_-pn2) placed in a separate drop and culture continued in KSOM. Where appropriate, embryos at the 2-cell stage were transferred to pseudopregnant CD-1 (ICR) females by as described ^52^. As a proof-of-principle, we generated a single offspring by cumulus cell nuclear transfer essentially as described previously ^56^.

### Preparation of ahES cells for nt

Following culture of FACS-purified ahES cells at 37°C in humidified 5% (v/v) CO_2_ in air, cell suspensions were prepared as previously described ^13,54^. Briefly, cells were washed with DMEM medium followed by calcium-free PBS and incubated with trypsin/EDTA for 3 min at 37°C. Trypsinization was quenched by the addition of 5 ml ES/DMEM (DMEM supplemented with 5% [v/v] FCS/LIF; Leeb and Wutz, 2011) and cells dissociated by gentle pipetting. Single cell suspensions were pelleted by centrifugation (1,100rpm, 5 min) and resuspended in fresh ES/DMEM medium. Single cell ahES cell suspensions were placed on ice and used immediately for micromanipulation. In some cases, haploid cells were enriched by FACS sorting immediately prior to micromanipulation. Cell aggregates were removed by passing suspensions though a 50 μm cell strainer (FALCON) into a polypropylene FACS tube (BD). To avoid Hoechst toxicity, we employed SSC and FSC as FACS Aria parameters for haploid and diploid population separation. Enriched haploid ES cells were collected into an ice-cold FACS tube containing 1 ml ES/DMEM supplemented with serum and immediately used for micromanipulation. G1 cell selection was further attempted by selecting smaller cells as nucleus donors.

### Differentiation assay

To evaluate differentiation potential of parthenogenetic and androgenetic cells, the expression level of naïve pluripotency and early differentiation markers was analysed in comparison to biparental control by RT-qPCR. ES-m1, ES-m2, ES-f1, pES1, pES3 pES4, aES1 and aES2 cell lines were plated in N2B27 based 2i/LIF medium at a final density of 10^4^ cells/cm^2^. On the next day, cells were washed with PBS and medium was exchanged to either N2B27 without 2i/LIF to induce differentiation, or fresh N2B27 + 2i/LIF for the undifferentiated controls. After 24h, cells were washed twice with PBS and harvested in RNA Lysis buffer + 1% 2-merchaptothanol and stored at −80°C before isolation of RNA using the EXTRACTME Total RNA Kit (Blirt). RNA was reverse-transcribed into cDNA using the SensiFAST cDNA Synthesis Kit (Bioline). Pluripotency and early differentiation marker as well as selected known imprinted genes expression was determined by qPCR using the Sensifast SYBR No Rox-Kit (Bioline).

### Statistical analysis

Statistical differences between pairs of data sets were analysed by a Chi-squared test.

### Imprinted gene assignment from RNA-seq data

Single blastocysts from reciprocal crosses (natural mating) between *Mus musculus musculus* C57BL/6 (B6) and *Mus musculus castaneus* (*cast*) were lysed and RNA extracted. Samples were processed using a SMART-Seq2 compatible protocol and processed as described ^25^. For imprinted gene assignment, we tested whether the difference between paternal and maternal reads per gene was significant using DESeq2. Only reads covering annotated SNPs for B6 and *cast* were used for the analysis. The number of reads per gene that could be assigned to one of the strains by the SNP information was taken from an intermediate result of the Allelome.Pro pipeline ^8,57^ and used to create a count table for all samples from forward (B6 x Cast) and reverse crosses (Cast x B6). Samples 4 (less than 5 % of reads mapping to the reference genome) and 6 (too low number of total reads) were removed from the analysis pipeline. Moreover, genes on the X-chromosome and genes with fewer than ten SNP spanning reads in at least one sample were removed from further analysis. We then employed DESeq2 to test for significant differences in maternal and paternal expression, in addition to delineating strain-specific expression using an adjusted *p*-value ≤ 0.05 as cut-off; genes that passed this threshold but were not consistently strain biased across all samples were not considered to be strain specific. This is the case e.g. for significantly regulated genes that are, however, detected only in one direction of crosses. The list of ‘published imprinted genes’ was comprises genes previously reported to be imprinted in the literature ^1,8^ and genes present in 4 different imprinting repositories (Mousebook, Otago, Geneimprint, Wamidex). From 388 unique imprinted gene names, 238 were also found in our dataset and could be assigned mgi gene symbols. Of these genes, 10 were located on the X chromosome and 51 were not represented by at least 10 SNP-overlapping reads; these genes were excluded from further analysis while 178 genes remained in the analysis pipeline. An additional 3 gene names were associated with predicted genes and hence removed from further analysis. To further exclude the possibility of erroneously calling imprinted genes due to reads assigned to overlapping transcripts, we only included genes if reads could be unambiguously assigned to a specific transcript. We further excluded genes that were not robustly expressed (genes with less than 12 reads in at least 4 samples). Genes with a L2FC > |0.5| between maternal and paternal alleles that fulfilled previous criteria and were consistently parentally biased across all samples were defined as ‘blastocyst-skewed expressed’ genes (BsX genes). Additionally, genes that showed a 70:30 expression ratio in at least 60% of samples in each cross between the parental alleles were considered as BiX genes (‘blastocyst-imprinted expressed’ genes).

### Evaluation of imprinted gene sets

We examined the identified and previously published imprinted genes using three bioinformatics approaches: First, gene set enrichment analysis was performed using Enrichr ^49^ against a database of genes whose disruption is associated with known phenotypic changes (MGI_Mammalian_Phenotype_2017) and against a database of genes deregulated in loss-of-function experiments (TF-LOF Expression_from_GEO). The top 8 terms sorted by p-value were selected. Second, we obtained allele-specific single-cell gene expression data from oocytes and preimplantation embryos ^18^ from GEO (GSE80810) and used these data to confirm parentally-biased allele-specific expression of published, nBiX, nBsX, but not of unconfirmed published imprinted genes throughout preimplantation development. Third, comparison of absolute log2FCs between ESCs cultured in 2i and 24h after induction of differentiation by 2i withdrawal ^24^ for the different groups of genes. Genes showing parentally-biased allele-specific expression, genes showing equal expression from both alleles, known imprinted genes and all genes were used as gene groups to compare the dynamics in gene expression.

### Whole genome bisulfite sequencing

Sequencing libraries for DNA methylation mapping were prepared using the μWGBS protocol^25^. Starting directly from lysed cells in digestion buffer, proteinase K digestion was performed at 50°C for 20 minutes. Custom-designed methylated and unmethylated oligonucleotides were added at a concentration of 0.1% to serve as spike-in controls for monitoring bisulfite conversion efficiency. Bisulfite conversion was performed using the EZ DNA Methylation-Direct Kit (Zymo Research, D5020) according to the manufacturer’s protocol, with the modification of eluting the DNA in only 9 μl of elution buffer. Bisulfite-converted DNA was used for single-stranded library preparation using the EpiGnome Methyl-Seq kit (Epicentre, EGMK81312) with the described modifications. Quality control of the final library was performed by measuring DNA concentrations using the Qubit dsDNA HS assay (Life Technologies, Q32851) on Qubit 2.0 Fluorometer (Life Technologies, Q32866) and by determining library fragment sizes with the Agilent High Sensitivity DNA Analysis kit (Agilent, 5067-4626) on Agilent 2100 Bioanalyzer Station (Agilent, G2939AA). All libraries were sequenced by the Biomedical Sequencing Facility at CeMM using the 2×75bp paired-end setup on the Illumina HiSeq 3000/4000 platform.

### DNA methylation data processing

Sequencing adapter fragments were trimmed using Trimmomatic v0.32 ^58^. The trimmed reads were aligned with Bismark v0.12.2 ^59^ with the following parameters: --*minins 0 --maxins 6000 --bowtie2*, which uses Bowtie2 v2.2.4 ^60^ for read alignment to the mm10 assembly of the mouse reference genome. Duplicate reads were removed as potential PCR artefacts and reads with a bisulfite conversion rate below 90% or with fewer than three cytosines outside a CpG context (required to confidently assess bisulfite conversion rate) were removed as potential post-bisulfite contamination. DNA methylation levels estimated by the Bismark extractor were loaded into R retaining all CpGs that were covered with at least three reads in at least two samples. We then used *dmrseq* ^31^ to identify consistently methylated regions of neighboring CpGs (*n*=168,061 regions) between androgenote, parthenogenote, and ICSI blastocysts (two replicates per sample group, total *n*=6). We retained all regions with opposing DNA methylation levels in uniparental vs. ICSI blastocysts (i.e., either *ah* > *ICSCI* > *ph*, or *ah* < *ICSI* < *ph*), with at least 100 reads total coverage (across all replicates), and with a minimum length of 100bp. Testing those regions (*n*=77,358) for significant differences in DNA methylation levels by sample group (FDR-adjusted p-value <= 0.1, |ah-ph| >= 30 percentage points, |β_ah_| >= 0.25, |β_ph_| >= 0.25) yielded 859 candidate DMRs. To enable comparison of these DMRs with the DNA methylation status in oocytes, sperm, and the ICM, we obtained published MethylC-Seq data (Wang et al., 2014) from GEO (GSE56697).

### Positional, region overlap, and motif enrichment analysis

We examined the identified genomic DMR regions with two complementary approaches: First, we used Locus Overlap Analysis ^33^ (LOLA; v1.12.0) to identify significant overlaps with experimentally determined transcription factor binding sites from publicly available ChIP-seq data. To this end, we used 791 ChIP-seq peak datasets from the LOLA Core database and we additionally added Znf445 binding peaks ^37^. We considered only terms with an 8-fold enrichment and an FDR-adjusted p-value below 0.005 significant. We focused on datasets from embryonic stem cells that were at least 1.5-fold more enriched in DMRs than in promoter regions. Second, we searched the DNA sequences underlying each DMR for matches to known DNA binding motifs from the HOCOMOCO database v11 ^38^. For this search, we used FIMO ^50^ (v4.10.2) (parameters: --*no-qvalue --text --bgfile motif-file*), and regions with at least one hit (p < 0.0001) were counted. To test for motif enrichment, we used Fisher’s exact test. Motifs with a 4-fold enrichment and an FDR-adjusted p-value below 0.005 were considered significant. We focused on motifs at least 1.5-fold more enriched in DMRs than in promoter regions.

### Analysis of allele-specific H3K27me3

We defined parent-of-origin-specific H3K27me3 imprints using published allele-specific ChIP-seq data from the ICM ^41^. To this end, we downloaded peak coordinates from GEO (GSE76687), converted the coordinates to the mm10 genome assembly using liftOver, and associated peaks with a gene if a peak was found within 5kb of its transcription start site.

### Analysis of DNA-methylation-dependent and H3K27me3-dependent allelic expression

To assess dependence of the allelic expression bias of imprinted genes on DNA methylation and H3K27me3, we obtained allele-specific RNA-seq data before and after maternal knockout (mKO) of *Dnmt3l* and *Eed* ^42,43^ from GEO (GSE130115 and GSE116713). First, genes were defined as imprinted in morulae if they showed a log2FC between maternal and paternal alleles > |1| in control morulae in at least one of the two datasets. Genes that lose their imprinted-like status (log2FC control/log2FC KO >1 & log2FC KO between −1 and 1) upon maternal deletion of *Dnmt3l* or *Eed* were considered to be dependent on maternally deposited DNA methylation or H3K27me3, respectively.

### E4.5 and E6.5 embryo RNA extraction, RT-PCR and Sanger Sequencing

RNA was extracted from single embryos and processed using a SMARTSeq2 compatible protocol. Resulting cDNA was used as a template to amplify PCR fragments covering at least one SNP per gene. Resulting fragments were then analysed by Sanger Sequencing.

### Topologically associating domains (TAD)

Because TAD positions are known to be relatively invariant across different tissues, we employed TADs coordinates defined in mouse ES cells ^61^ as a proxy for E4.5 blastocysts. DMR and imprinted genes are defined to be within the same TAD if the centre of DMR and gene transcription start sites are located within the same TADs (DMRs falling into TAD boarders up to 50kb are excluded from downstream analysis). The control set was generated by randomly shifting genome coordinates of both genes and DMR on each chromosome 1000 times, so that pair-wise distances between genes and DMR are kept the same. The number of imprinted genes having a DMR associated within the same TAD was calculated for both the control and the experiment datasets.

## REFERENCES

1. Tucci, V., Isles, A.R., Kelsey, G., Ferguson-Smith, A.C. & Erice Imprinting, G. Genomic Imprinting and Physiological Processes in Mammals. Cell 176, 952–965 (2019).

2. Sanli, I. & Feil, R. Chromatin mechanisms in the developmental control of imprinted gene expression. The International Journal of Biochemistry & Cell Biology 67, 139–147 (2015).

3. Surani, M.A., Barton, S.C. & Norris, M.L. Development of reconstituted mouse eggs suggests imprinting of the genome during gametogenesis. Nature 308, 548–50 (1984).

4. McGrath, J. & Solter, D. Completion of mouse embryogenesis requires both the maternal and paternal genomes. Cell 37, 179–83 (1984).

5. Morison, I.M., Ramsay, J.P. & Spencer, H.G. A census of mammalian imprinting. Trends Genet 21, 457–65 (2005).

6. Schulz, R. et al. WAMIDEX: a web atlas of murine genomic imprinting and differential expression. Epigenetics 3, 89–96 (2008).

7. Blake, A. et al. MouseBook: an integrated portal of mouse resources. Nucleic Acids Res 38, D593–9 (2010).

8. Andergassen, D. et al. Mapping the mouse Allelome reveals tissue-specific regulation of allelic expression. Elife 6 (2017).

9. Inoue, A., Jiang, L., Lu, F., Suzuki, T. & Zhang, Y. Maternal H3K27me3 controls DNA methylation-independent imprinting. Nature 547, 419 (2017).

10. Bourc’his, D., Xu, G.L., Lin, C.S., Bollman, B. & Bestor, T.H. Dnmt3L and the establishment of maternal genomic imprints. Science 294, 2536–9 (2001).

11. Schulz, R. et al. The parental non-equivalence of imprinting control regions during mammalian development and evolution. PLoS Genet 6, e1001214 (2010).

12. Ferguson-Smith, A.C. & Bourc’his, D. The discovery and importance of genomic imprinting. Elife 7 (2018).

13. Leeb, M. & Wutz, A. Derivation of haploid embryonic stem cells from mouse embryos. Nature 479, 131–4 (2011).

14. Leeb, M. et al. Germline potential of parthenogenetic haploid mouse embryonic stem cells. Development 139, 3301–5 (2012).

15. Yang, H. et al. Generation of genetically modified mice by oocyte injection of androgenetic haploid embryonic stem cells. Cell 149, 605–17 (2012).

16. Wan, H. et al. Parthenogenetic haploid embryonic stem cells produce fertile mice. Cell Research 23, 1330–1333 (2013).

17. Li, Z.-K. et al. Generation of Bimaternal and Bipaternal Mice from Hypomethylated Haploid ESCs with Imprinting Region Deletions. Cell Stem Cell 23, 665–676.e4 (2018).

18. Borensztein, M. et al. Xist-dependent imprinted X inactivation and the early developmental consequences of its failure. Nat Struct Mol Biol 24, 226–233 (2017).

19. Monk, D., Mackay, D.J.G., Eggermann, T., Maher, E.R. & Riccio, A. Genomic imprinting disorders: lessons on how genome, epigenome and environment interact. Nature Reviews Genetics 20, 235–248 (2019).

20. Duffié, R. et al. The Gpr1/Zdbf2 locus provides new paradigms for transient and dynamic genomic imprinting in mammals. Genes & Development 28, 463–478 (2014).

21. Czermin, B. et al. <em>Drosophila</em> Enhancer of Zeste/ESC Complexes Have a Histone H3 Methyltransferase Activity that Marks Chromosomal Polycomb Sites. Cell 111, 185–196 (2002).

22. Terranova, R. et al. Polycomb Group Proteins Ezh2 and Rnf2 Direct Genomic Contraction and Imprinted Repression in Early Mouse Embryos. Developmental Cell 15, 668–679 (2008).

23. Pereira, C.F. et al. ESCs require PRC2 to direct the successful reprogramming of differentiated cells toward pluripotency. Cell Stem Cell 6, 547–56 (2010).

24. Lackner, A. et al. Cooperative molecular networks drive a mammalian cell state transition. bioRxiv, 2020.03.23.000109 (2020).

25. Farlik, M. et al. Single-cell DNA methylome sequencing and bioinformatic inference of epigenomic cell-state dynamics. Cell Rep 10, 1386–97 (2015).

26. Li, W. et al. Androgenetic haploid embryonic stem cells produce live transgenic mice. Nature 490, 407–11 (2012).

27. Wang, L. et al. Programming and Inheritance of Parental DNA Methylomes in Mammals. Cell 157, 979–991 (2014).

28. Gigante, S. et al. Using long-read sequencing to detect imprinted DNA methylation. Nucleic Acids Research 47, e46–e46 (2019).

29. Matsuzaki, H., Okamura, E., Shimotsuma, M., Fukamizu, A. & Tanimoto, K. A Randomly Integrated Transgenic <em>H19</em> Imprinting Control Region Acquires Methylation Imprinting Independently of Its Establishment in Germ Cells. Molecular and Cellular Biology 29, 4595–4603 (2009).

30. Li, X. et al. A Maternal-Zygotic Effect Gene, <em>Zfp57</em>, Maintains Both Maternal and Paternal Imprints. Developmental Cell 15, 547–557 (2008).

31. Korthauer, K., Chakraborty, S., Benjamini, Y. & Irizarry, R.A. Detection and accurate false discovery rate control of differentially methylated regions from whole genome bisulfite sequencing. Biostatistics 20, 367–383 (2018).

32. Braidotti, G. et al. The Air noncoding RNA: an imprinted cis-silencing transcript. Cold Spring Harb Symp Quant Biol 69, 55–66 (2004).

33. Sheffield, N.C. & Bock, C. LOLA: enrichment analysis for genomic region sets and regulatory elements in R and Bioconductor. Bioinformatics 32, 587–589 (2015).

34. Sánchez-Castillo, M. et al. CODEX: a next-generation sequencing experiment database for the haematopoietic and embryonic stem cell communities. Nucleic Acids Research 43, D1117–D1123 (2014).

35. Dunham, I. et al. An integrated encyclopedia of DNA elements in the human genome. Nature 489, 57 (2012).

36. Leung, D. et al. Regulation of DNA methylation turnover at LTR retrotransposons and imprinted loci by the histone methyltransferase Setdb1. Proceedings of the National Academy of Sciences 111, 6690–6695 (2014).

37. Takahashi, N. et al. ZNF445 is a primary regulator of genomic imprinting. Genes Dev 33, 49–54 (2019).

38. Kulakovskiy, I.V. et al. HOCOMOCO: towards a complete collection of transcription factor binding models for human and mouse via large-scale ChIP-Seq analysis. Nucleic Acids Research 46, D252–D259 (2017).

39. Rodriguez-Jato, S., Nicholls, R.D., Driscoll, D.J. & Yang, T.P. Characterization of cis- and trans-acting elements in the imprinted human SNURF-SNRPN locus. Nucleic Acids Res 33, 4740–53 (2005).

40. Dixon, J.R. et al. Topological domains in mammalian genomes identified by analysis of chromatin interactions. Nature 485, 376–380 (2012).

41. Zheng, H. et al. Resetting Epigenetic Memory by Reprogramming of Histone Modifications in Mammals. Molecular Cell 63, 1066–1079 (2016).

42. Chen, Z., Yin, Q., Inoue, A., Zhang, C. & Zhang, Y. Allelic H3K27me3 to allelic DNA methylation switch maintains noncanonical imprinting in extraembryonic cells. Science Advances 5, eaay7246 (2019).

43. Inoue, A., Chen, Z., Yin, Q. & Zhang, Y. Maternal Eed knockout causes loss of H3K27me3 imprinting and random X inactivation in the extraembryonic cells. Genes Dev 32, 1525–1536 (2018).

44. Ferguson-Smith, A.C. Genomic imprinting: the emergence of an epigenetic paradigm. Nature Reviews Genetics 12, 565–575 (2011).

45. Verona, R.I., Mann, M.R. & Bartolomei, M.S. Genomic imprinting: intricacies of epigenetic regulation in clusters. Annu Rev Cell Dev Biol 19, 237–59 (2003).

46. Barlow, D.P. & Bartolomei, M.S. Genomic Imprinting in Mammals. Cold Spring Harbor Perspectives in Biology 6 (2014).

47. Choi, J. et al. Prolonged Mek1/2 suppression impairs the developmental potential of embryonic stem cells. Nature 548, 219–223 (2017).

48. Sagi, I. et al. Distinct Imprinting Signatures and Biased Differentiation of Human Androgenetic and Parthenogenetic Embryonic Stem Cells. Cell Stem Cell 25, 419–432 e9 (2019).

49. Chen, E.Y. et al. Enrichr: interactive and collaborative HTML5 gene list enrichment analysis tool. BMC Bioinformatics 14, 128 (2013).

50. Grant, C.E., Bailey, T.L. & Noble, W.S. FIMO: scanning for occurrences of a given motif. Bioinformatics 27, 1017–1018 (2011).

51. Yoshida, N. & Perry, A.C. Piezo-actuated mouse intracytoplasmic sperm injection (ICSI). Nat Protoc 2, 296–304 (2007).

52. Suzuki, T., Asami, M. & Perry, A.C. Asymmetric parental genome engineering by Cas9 during mouse meiotic exit. Sci Rep 4, 7621 (2014).

53. Suzuki, T. et al. Mice produced by mitotic reprogramming of sperm injected into haploid parthenogenotes. Nature Communications 7, 12676 (2016).

54. Leeb, M., Perry, A.C.F. & Wutz, A. Establishment and Use of Mouse Haploid ES Cells. Curr Protoc Mouse Biol 5, 155–185 (2015).

55. Leeb, M., Dietmann, S., Paramor, M., Niwa, H. & Smith, A. Genetic exploration of the exit from self-renewal using haploid embryonic stem cells. Cell Stem Cell 14, 385–93 (2014).

56. Wakayama, T., Perry, A.C.F., Zuccotti, M., Johnson, K.R. & Yanagimachi, R. Full-term development of mice from enucleated oocytes injected with cumulus cell nuclei. Nature 394, 369–374 (1998).

57. Andergassen, D. et al. Allelome.PRO, a pipeline to define allele-specific genomic features from high-throughput sequencing data. Nucleic Acids Res 43, e146 (2015).

58. Bolger, A.M., Lohse, M. & Usadel, B. Trimmomatic: a flexible trimmer for Illumina sequence data. Bioinformatics 30, 2114–2120 (2014).

59. Krueger, F. & Andrews, S.R. Bismark: a flexible aligner and methylation caller for Bisulfite-Seq applications. Bioinformatics 27, 1571–1572 (2011).

60. Langmead, B. & Salzberg, S.L. Fast gapped-read alignment with Bowtie 2. Nature Methods 9, 357–359 (2012).

61. Stevens, T.J. et al. 3D structures of individual mammalian genomes studied by single-cell Hi-C. Nature 544, 59–64 (2017).

